# RTEL1 is required for silencing and epigenome stability

**DOI:** 10.1101/2023.04.11.535823

**Authors:** Margaux Olivier, Amy Hesketh, Marie-Noëlle Pouch-Pélissier, Thierry Pélissier, Ying Huang, David Latrasse, Moussa Benhamed, Olivier Mathieu

## Abstract

Transcriptional silencing is an essential mechanism for controlling the expression of genes, transgenes and heterochromatic repeats through specific epigenetic marks on chromatin that are maintained during DNA replication. In Arabidopsis, silenced transgenes and heterochromatic sequences are typically associated with high levels of DNA methylation, while silenced genes are enriched in H3K27me3. Reactivation of these loci is often correlated with decreased levels of these repressive epigenetic marks. Here, we report that the DNA helicase REGULATOR OF TELOMERE ELONGATION 1 (RTEL1) is required for transcriptional silencing. RTEL1 deficiency causes upregulation of many genes enriched in H3K27me3 accompanied by a moderate decrease in this mark, but no loss of DNA methylation at reactivated heterochromatic loci. Instead, heterochromatin exhibits DNA hypermethylation and increased H3K27me3 in *rtel1*. We further find that loss of RTEL1 suppresses the release of heterochromatin silencing caused by the absence of the MOM1 silencing factor. RTEL1 is conserved among eukaryotes and plays a key role in resolving DNA secondary structures during DNA replication. Inducing such aberrant DNA structures using DNA cross-linking agents also results in a loss of transcriptional silencing. These findings uncover unappreciated roles for RTEL1 in transcriptional silencing and in stabilizing DNA methylation and H3K27me3 patterns.

## Introduction

In eukaryotic cells, DNA is packaged with histone proteins to form a highly compacted and organized structure known as chromatin. Chromatin exists in two different main states: euchromatin, which is mainly composed of protein coding genes and sparsely condensed, and heterochromatin, which shows a high density of transposable elements and repetitive sequences and a higher level of condensation. By influencing DNA accessibility, chromatin structure largely influences various DNA-related biological processes, including transcription, DNA replication and DNA repair. The chromatin structure is also related to the presence of epigenetic marks, including various post-translational modifications of histone proteins and DNA methylation.

In *Arabidopsis thaliana,* DNA methylation of cytosine residues occurs in three sequence contexts: the symmetric CG and CHG contexts and the asymmetric CHH context (where H is any base but G) (1). Typically, heterochromatin sequences are transcriptionally silent and associated with high levels of DNA methylation at all three cytosine sequence contexts. In contrast, most constitutively active genes in euchromatin are either unmethylated or DNA methylated in their coding sequence specifically at CG sites (2–4). On the other hand, genes with spatiotemporal expression are typically silenced in certain tissues or at certain developmental stages by histone H3 lysine 27 trimethylation (H3K27me3) deposited by Polycomb group (PcG) proteins (5, 6).

DNA methylation is established *de novo* by the RNA-directed DNA methylation (RdDM) pathway and then maintained by specific pathways to each cytosine sequence context. CG sites are the most frequently and highly methylated, with maintenance dependent on METHYLTRANSFERASE 1 (MET1) that acts during DNA replication. CHROMOMETHYLASE 3 (CMT3) is the primary methyltransferase for CHG methylation maintenance. Methylation at CHH sites in heterochromatin is mainly maintained by CHROMOMETHYLASE 2 (CMT2), while DOMAINS REARRANGED METHYLTRANSFERASE 2 (DRM2) acting in the RdDM pathway methylates CHH positions in shorter, less heterochromatic sequences (1, 7). In the poorly accessible heterochromatin environment, DNA methylation depends on Decrease in DNA Methylation 1 (DDM1) to overcome the linker histone H1 barrier and allow access to DNA for MET1, CMT2 and CMT3 (8, 9).

Disruption of DNA methylation pathways, particularly CG methylation, leads to the reactivation of numerous transposable elements and some protein coding genes (3, 4, 10). However, DNA methylation is not the sole mechanism which the cell relies on to secure transcriptional gene silencing. There is growing evidence that DNA methylation-independent pathways participate in robust silencing. Several key factors involved in these pathways have been identified, including MORPHEUS’ MOLECULE 1 (MOM1), a protein that mediates silencing of various pericentromeric sequences (11–14), and members of the conserved *Microrchidia (MORC)* ATPase family, AtMORC1 and AtMORC6, which regulate silencing of several, mostly heterochromatic, TEs (15). The precise mechanisms by which these actors control silencing are not yet fully understood. There is evidence that MOM1 and AtMORC6 act in largely distinct pathways in controlling silencing of pericentromeric TEs, although MOM1 appears to recruit the RdDM machinery via MORC6 at some RdDM sites (16, 17). Transcriptional reactivation in *mom1* mutants requires the MED14 subunit of the MEDIATOR complex (18). More recently, we found that the MAINTENANCE OF MERISTEMS (MAIN) and MAIN-LIKE1 (MAIL1) proteins, which contain a transposon-related plant mobile domain, also control silencing of a set of pericentromeric TEs independently of DNA methylation through a molecular pathway distinct from those involving MOM1 and AtMORC6 (19).

During DNA replication, both heterochromatin and euchromatin are disrupted ahead of the replication fork and are then reassembled into their original epigenetic states behind the fork (20). Similarly, chromatin structure must be modulated during DNA repair to allow repair factors access to the DNA molecule, and subsequently restored to its original state to maintain transcriptional activity of the underlying sequences. Stalling of the replication fork leads to epigenetic instability and loss of silencing, suggesting that DNA unwinding, replication and chromatin re-formation must be tightly coordinated (21). Mutations in several factors involved in DNA replication and/or repair have been found to affect silencing, indicating a tight link between these processes. These factors include, among others, the DNA polymerases POLα (22), POLδ (23), POL (24, 25), REPLICATION PROTEIN 2 (RPA2) (26), the nuclear protein BRUSHY1 (BRU1) (27), the flap endonuclease FEN1 (28) and the chromatin assembly factor CAF-1 (29). Mutants with defects in these factors exhibit increased sensitivity to DNA damaging agents, indicating the accumulation of DNA damage and/or constitutive replication stress. Notably, release of silencing in these mutants is not associated with a loss of DNA methylation.

Here, we identify a role in silencing for *REGULATOR OF TELOMERE ELONGATION 1* (*RTEL1*) encoding a DNA helicase protein linked to DNA repair and replication. We show that RTEL1 is required for the silencing of both a transgene and a set of endogenous loci independently of a loss of DNA methylation, and through a pathway likely related to its replication function. Additionally, we show that RTEL1 is necessary from preventing a gain of H3K27me3 and DNA methylation in heterochromatin regions. Upregulated genes in *rtel1* show a moderate loss of H3K27me3, while heterochromatin regions gain H3K27me3 and DNA methylation. Moreover, our study indicates that RTEL1 is required for the release of heterochromatin silencing in the absence of MOM1, and for viably in *pol2a* and *mail1* mutant backgrounds.

## Materials and Methods

### Plant material

Plants were grown in soil in long-day conditions (16 h light, 8 h dark) at 23 °C with 60% relative humidity. The *rtel1-3* mutant allele reported in this study was isolated from a population of EMS mutagenized L5 plants that we previously described (19). The *rtel1-3* mutant was backcrossed twice to the parental L5 line before analysis. The *rtel1-1* (SALK_113285), *pol2a-12* (24)*, mail1-1* (GK-840E05), *mom1-2* (SAIL_610_G01) and *fancjb-1* (SALK_101493) mutant lines used in this study were all in a L5 genetic background. Primers used in this study are listed in Supplementary Table S6.

### Histochemical staining

Whole seedlings or rosette leaves were vaccum infiltrated twice 5 min with 3 ml of X- Gluc staining solution (50 mM NaHPO4 pH 7; 10mM EDTA; 0.2% Triton-X-100; 0.04% X-Gluc) and incubated 24 h at 37°C in the dark. Chlorophyll was subsequently cleared with repeated washes in ethanol at room temperature.

### Mutagenesis and mapping

The mutant L5 population used in this study was obtained by mutagenesis as described in Ikeda *et al.*, 2017. To screen for mutants impaired in transcriptional gene silencing, one leaf per M2 plant was dissected and histochemically stained as previously described. DNA of plant selected at this stage was extracted from rosette leaves using Wizard Genomic DNA Purification kit (Promega) and digested with methylation-sensitive restriction enzyme (MspI or CfoI, NEB) according to the manufacturer’s instructions. GUS 5’ extremity and FWA gene (control region without MspI/CfoI restriction site) was amplified, and methylation levels were estimated by measuring the amplification ratio between the transgene and the control gene to select mutants that reactivate L5-GUS transgene independently of a loss of DNA methylation. We performed mapping-by-sequencing using genomic DNA from a pool of 50-100 individual F2 mutant plants segregating from a *p14 12-1* x Ler cross. Libraries were prepared using the TruSeq Nano DNA Sample Preparation kit (Illumina) and sequenced on a NextSeq 500 instrument (Illumina) at Fasteris S.A. (Geneva, Switzerland) to generate ∼ 100M 150-bp paired-end reads. Reads were mapped on the *Arabidopsis* TAIR10 genome using BWA (30) with the default parameters. Single-nucleotide polymorphisms calling was performed using SAMtools mpileup followed by BCFtools view with the -bvgN option. Three candidate genes with EMS-induced non-synonymous mutations were identified in the mapping interval on chromosome 1. T-DNA mutant lines for the three candidate genes were analyzed for release of gene silencing and allelic tests were performed, allowing identification of *RTEL1*.

### Complementation of *rtel1-3*

A genomic fragment containing the *RTEL1* gene with its promoter region (1 kb upstream the ATG codon) was PCR amplified from Col-0 genomic DNA and cloned into the pDONR221 vector (Invitrogen) by BP recombination. Following sequence verification, this fragment was introduced in the GATEWAY destination vector pB7FWG2,0 Δp35S by LR recombination. The resulting *pRTEL1::RTEL1* construct was introduced into *rtel1-3* plants using *Agrobacterium-*mediated floral-dip transformation.

### Drug Treatments

The seeds were surface-sterilized with 0.05% SDS, 70% ethanol for 5 min, rinsed 5 min with 95% ethanol, air dried and evenly spread on sterile 0.8% agar, 1% sucrose and 1X Murashige and Skoog (M0255; Duchefa Biochemie) medium supplemented with or without the indicated concentrations of MMC (M0503; Sigma-Aldrich) or CP (TEVA Pharma). Whole seedlings were harvest at 14-day-old for analysis.

### Transcript analysis

Total RNA was extracted from immature inflorescences (stages 1-12) or aerial parts of 16- day-old seedlings using NucleoSpin RNA Plus kit (Macherey-Nagel) following the manufacturer’s recommendations. Gel-based RT-PCR analyses were performed using the One-Step RT-PCR kit (Qiagen) in a final volume of 10 µL starting from 0.03 µg of DNase-treated (RQ1 Rnase-free Dnase supplemented with Rnasin Ribonuclease Inhibitor; Promega) total RNA. For RT-qPCR analyses, 1 µg of Dnase-treated total RNA was reverse transcribed using the M-MLV Reverse Transcriptase (Promega) and Random Hexadeoxynucleotides (Promega) in a final volume of 25 µL. Two microliters of cDNA was used for subsequent amplification using SensiFAST SYBR No-Rox Kit (Bioline) on an Eco™ Real-Time PCR System (Ilumina), following a program of 5 min at 95°C, 40 cycles of 10 s at 95°C, and 30 s at 60°C, in a final reaction volume of 10 µL. Amplification of *ACTIN2* gene transcripts was used as a reference for normalization and data were analyzed according to the 2-ΔΔCt method (31).

### mRNA sequencing

Total RNA was extracted from 14- or 16-day-old seedlings with RNAzol (Euromedex) and then treated with the RQ1 RNase-free DNase (Promega) supplemented with RNasin Ribonuclease Inhibitor (Promega), cleaned up either with phenol-chloroform and precipitated with ethanol or with RNA Clean & Concentrator -5 kit (Zymo Research). Two biological replicates were collected for each genotype. Libraries were prepared using the TruSeq Stranded mRNA Library Prep kit (Illumina) and sequenced on a HiSeq 4000 instrument (Illumina) at Fasteris S.A. (Geneva, Switzerland) to generate ∼ 26–35M 50-bp single-end reads. Reads were aligned to the TAIR10 Arabidopsis genome plus the L5-GUS transgene with STAR version 2.7.9a (32) retaining multi-mapped reads mapping up to 10 positions. Subsequent read counting on TEs and PCGs was performed with featureCounts version 2.0.3 (33) using the Araport11 genome annotation. Normalization and differential analyses were done using DESeq2 version 1.26.0 (34) with default parameters. Only loci with absolute fold-change ≥ 1 and Benjamini-Hochberg adjusted P values < 0.05 were considered differentially expressed. We also re-analyzed publicly available data for *mail1* (ERR1593753, ERR1593754, ERR1593761, ERR1593762 (19)), *pol2a-12* (ERR4194076 - ERR4194079 (24)) and *clf swn* (SRR1931614, SRR1931620, SRR2500947, SRR2962411 (35)).

### Gene Ontology analysis

GO enrichment analysis was performed using the PANTHER Overrepresentation Test (version 17.0; http://pantherdb.org/ (36–38)). Statistical significance was calculated by Fisher’s exact test and adjusted using the Bonferroni correction for multiple testing (multiplication of the single-test *P* value by the number of independent tests to obtain an expected error rate) and *P* values < 0.05 were deemed statistically significant. Fold enrichment corresponds to the ratio of upregulated PCGs numbers annotated in the ontology term over all genes annotated in this term and was considered overrepresented if it was greater than 1.

### Bisulfite sequencing (BS-seq)

Genomic DNA was extracted from 16-day-old seedlings with the Wizard Genomic DNA Purification Kit (Promega), following the manufacturer’s instructions. Sodium bisulfite conversion, library preparation, and sequencing on a HiseqXten were performed at the Beijing Genomics Institute (Hong Kong), producing ∼ 26–35M 150-bp paired-end reads. Data analysis was done using the Methylpy analysis pipeline v1.4.6 (39). Reads were filtered to remove PCR duplicates using Picard v2.23.3 and mapped to TAIR10 using Bowtie2 v2.4.1 (40). Only cytosines with a minimum coverage of 6 reads were retained. To calculate average methylation levels at specific regions, we used the add-methylation-level function from Methylpy (39) . TEs located in euchromatic arms and TEs located in pericentromeric regions were previously defined (41). We also re-analyzed publicly available data for *mail1* (ERR1593765 - ERR1593768 (19)) and *pol2a-12* (ERR4194158, ERR4194163 (24)).

### Nuclei isolation and microscopy

Rosette leaves from 3-week-old plants were fixed for one hour in 4% formaldehyde in 1X PBS and subsequently chopped on a glass slide. Tissues were covered with a coverslip, manually squashed and frozen in liquid nitrogen before removing the coverslip. Slides were washed twice in 1X PBS and DNA was stained with DAPI in Vectashield mounting medium (Vector Laboratories). Nuclei were visualized using a Zeiss Axio Imager Z.1 epifluorescence microscope (Carl Zeiss AG) with a PL Apochromat 100X/1.40 oil objective and images were captured with a Zeiss AxioCam MRm camera using the Zeiss ZEN software. The relative heterochromatin fraction was computed for each nucleus by calculating the ratio of the sum of chromocenter volume (area x intensity) over that of the entire nucleus using the ImageJ software.

### Chromatin immunoprecipitation followed by sequencing (ChIP-seq)

ChIP-seq was performed starting from 2-week-old seedlings grown in soil. Briefly, 1 g. of aerial tissues were fixed in 1% formaldehyde-containing MC buffer (42). Fixed tissues were nitrogen frozen and ground to fine powder by vortexing in the presence of seven ceramic beads without thawing the extracts. Nuclei were isolated as previously described (43) and resuspended in 1ml of 20mM Tris pH8, 2mM EDTA, 0.1% SDS 0.1%, 1X complete protease inhibitor cocktail (Roche). Chromatin was then sonicated in a Covaris S220 with the following settings: treatment time 5 min, acoustic duty factor 20%, PIP 170W, Cycles per burst 200 and max temperature 8°C. Following centrifugation at 13000rpm 4°C for 10 min., about half of the chromatin extract was used for immunoprecipitation with 5µg of H3K27me3 antibody (C15410069 Diagenode). IP, washes, and reverse crosslink were carried out as described in (42) . Reverse crosslinked DNA was purified using Zymo ChIP DNA clean and concentrator columns following the manufacturer’s instructions. Library preparation from the eluted DNA and PE100 sequencing on a DNBseq sequencing platform was performed at the Beijing Genomics institute (Hong Kong). Two biological replicates were performed for each genotype.

Reads were trimmed and filtered for quality and adapter contamination using Trim Galore (https://www.bioinformatics.babraham.ac.uk/projects/trim_galore/), and aligned to the TAIR10 genome plus the L5-GUS transgene using bowtie2. Multi-mapping reads were discarded, and probable PCR duplicates were removed using the Picard Tools suite. Sample tracks and metaplots over genomic regions were obtained using deeptools bamCoverage (- normalizeUsing RPGC).

### ATAC-seq

One hundred mg of 14-day-old seedlings were ground and nuclei were isolated with 4°C Buffer (0,25M Sucrose, 10mM Tris-HCl, 10mM MgCL2, 1% Triton, 5mM ß- mercaptoethanol) containing proteinase inhibitor cocktail (Roche) and filtered in 63 µm. Nuclei were resuspended in 1X TD Buffer (Illumina FC-121-1030) and 2.5 µl of Tn5 Transposase (Illumina FC-121-1030) were added. Tagmentation reaction was performed at 37°C for 30min, and DNA was purified using a Qiagen MinElute Kit. DNA libraries were amplified for a total of 8 cycles. The libraries were then subjected to 1×75bp high-throughput sequencing using a NextSeq 500 instrument (Illumina). Reads were mapped to the *Arabidopsis* TAIR10 reference genome using STAR version 2.7.2b (32). Peaks were called with MACS2 v2.2 (44). Data for generating metaplots was obtained with DeepTools v3.3.2 (45).

### Flow cytometry

Flow cytometry analysis was performed as previously described (19). Briefly, nuclei were prepared from 18-day-old seedlings using the CyStain UV Precise P kit (Partec), according to the manufacturer’s recommendations. Flow cytometry profiles were obtained on an Attune Acoustic Focusing Cytometer (Applied Biosystems) and analyzed with the Attune Cytometric software (Applied Biosystems).

## Results

### Mutations in *RTEL1* release transcriptional silencing

The *Arabidopsis* L5 line carries transcriptionally silent tandem repeats of a *ß-glucuronidase* (*GUS*) transgene under control of the CaMV 35S promoter (46). We and others previously showed that silencing of the L5-GUS transgene is sensitive to mutations in a wide range of transcriptional silencing regulators (18, 19, 24, 26, 27, 46). We conducted a genetic screen on an ethyl methanesulfonate (EMS) mutagenized L5 population for mutants that reactivate the L5-GUS transgene independently of a loss of DNA methylation. We used GUS histochemical staining assay on isolated leaves and methylation-sensitive restriction enzyme digestion followed by PCR to select M2 plants with destabilized L5-GUS silencing and no loss of DNA methylation at the 5’ end of the transgene GUS coding region. We identified six mutants, for which silencing release of the L5-GUS transgene followed a 1/3 ratio in the F2 progeny of backcrosses with L5, indicating that the causative mutations were single-locus nuclear recessive. Further mapping-by-sequencing of candidate mutations using outcross F2 populations identified new mutant alleles of *RPA2, MOM1, POLa2,* and two mutant alleles of *POL2A* already described (Supplementary Figure S1) (24, 25), validating our screening strategy as mutations in these genes are known to affect silencing without associated loss of DNA methylation (13, 22, 24–26). In the last mutant, *p14 12-1*, we identified a C to T substitution in *AT1G79950 (REGULATOR OF TELOMERE LENGTH 1, RTEL1*), causing a premature stop codon at the amino acid position 712 (Figure 1a-c, Supplementary Figure S2a), most likely leading to an abortion of translation. Complementation test between *p14 12- 1* and the independent *rtel1-1* T-DNA insertion mutant line (47), rescue of *p14 12-1* defects with a *RTEL1* genomic copy, and introducing L5-GUS in the *rtel1-1* background confirmed that mutation of *RTEL1* was responsible for the observed silencing defects (Figure 1, Supplementary Figure S2b-d). Hence, *p14 12-1* was renamed *rtel1-3*. In addition to the L5- GUS transgene, silencing of the Athila retrotransposon-derivates *TSI* repeats was destabilized to some extent in *rtel1* mutants, and expression of the silenced 45S rRNA variant VAR1 was activated in *rtel1-3* (Figure 1d).

**Figure 1:**
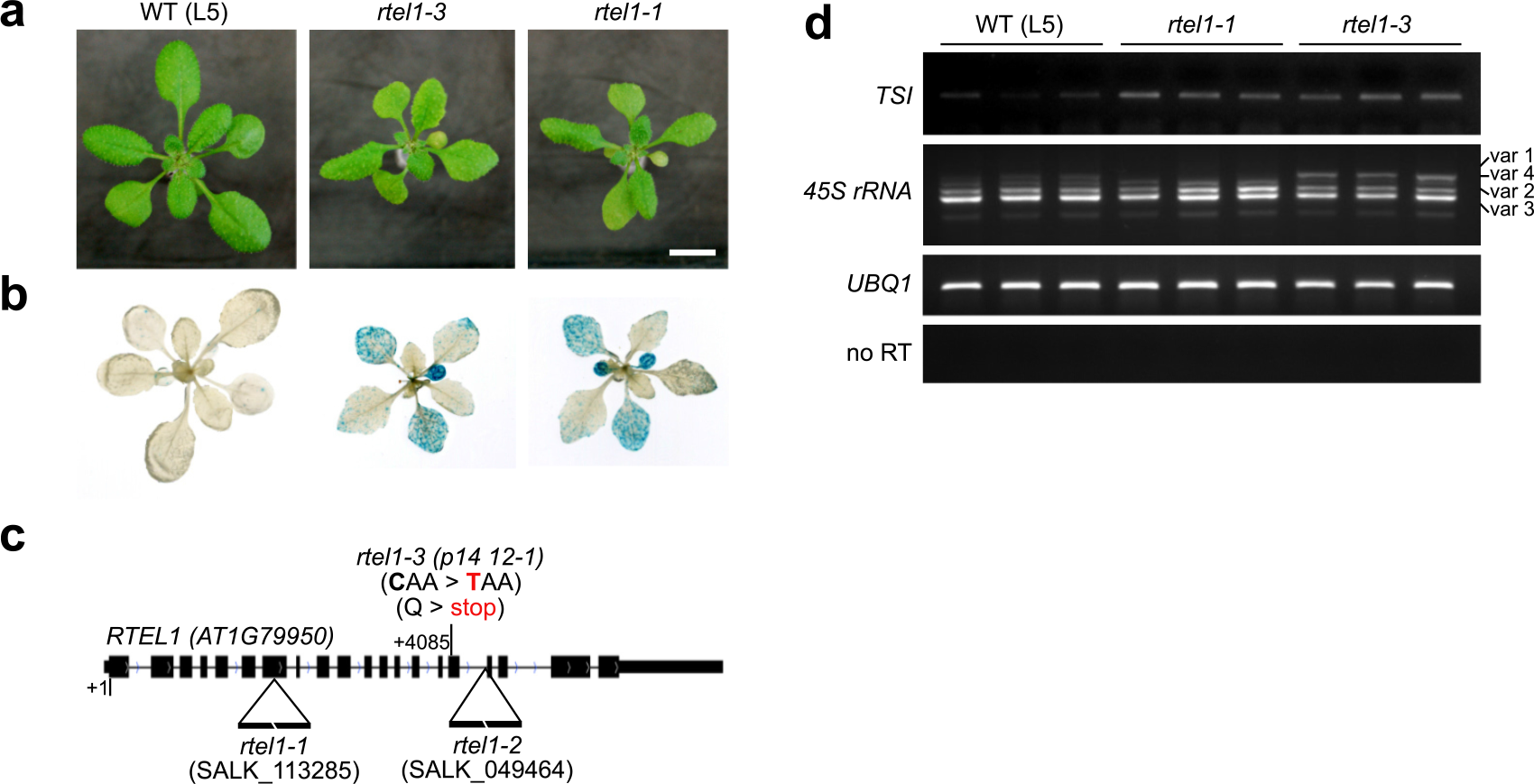
Mutations in *RTEL1* release transcriptional silencing of the L5-GUS transgene. (a) Photos of 3-week-old plants of the indicated genotypes. Scale bar, 1cm. **(b)** L5-GUS transgene activity detected by histochemical staining in 3-week-old plants of the indicated genotypes. **(c)** Representation of *RTEL1* gene with the insertion and mutation positions of the three mutant alleles. The point mutation identified in *rtel1-3* and the resulting amino acid change are indicated in red. **(d)** RT-PCR analysis of transcripts from endogenous repeats, *TSI* (*Transcriptionally Silent Information*) and 45S rRNA genes. Amplification of *UBQ1* (*UBIQUITIN EXTENSION PROTEIN 1*) was used as a loading control. For each target, PCR in the absence of reverse transcription (no RT) was performed to control for genomic DNA contamination.

### Loss of *rtel1* upregulates DNA damage response genes and genes enriched in H3K27me3

To define how mutations in *RTEL1* affect transcription across the genome, we conducted RNA sequencing (RNA-seq) on *rtel1-1* and *rtel1-3* mutant seedlings and compared their transcriptomes to that of WT (L5) plants. We identified 346 and 781 differentially expressed genes (DEGs) in *rtel1-1* and *rtel1-3,* respectively. The large majority of DEGs in *rtel1-1* (75%) and *rtel1-3* (90%) were upregulated, indicating that loss of *RTEL1* primarily leads to the upregulation of transcripts (Figure 2a). Notably, the upregulated transcripts were mostly derived from protein coding genes (PCGs), although 16 and five TEs were also upregulated in *rtel1-3* and *rtel1-1,* respectively. We identified 687 PCGs upregulated in *rtel1-3* and 261 in *rtel1-1*, with about 76% of the *rtel1-1* upregulated PCGs being also upregulated in *rtel1-3* (Figure 2b). Thus, *rtel1-3* has a stronger impact on transcription compared to *rtel1-1*. These results indicate that RTEL1 is required for proper expression of several hundreds of genes and silencing of a subset of heterochromatic repeats.

**Figure 2:**
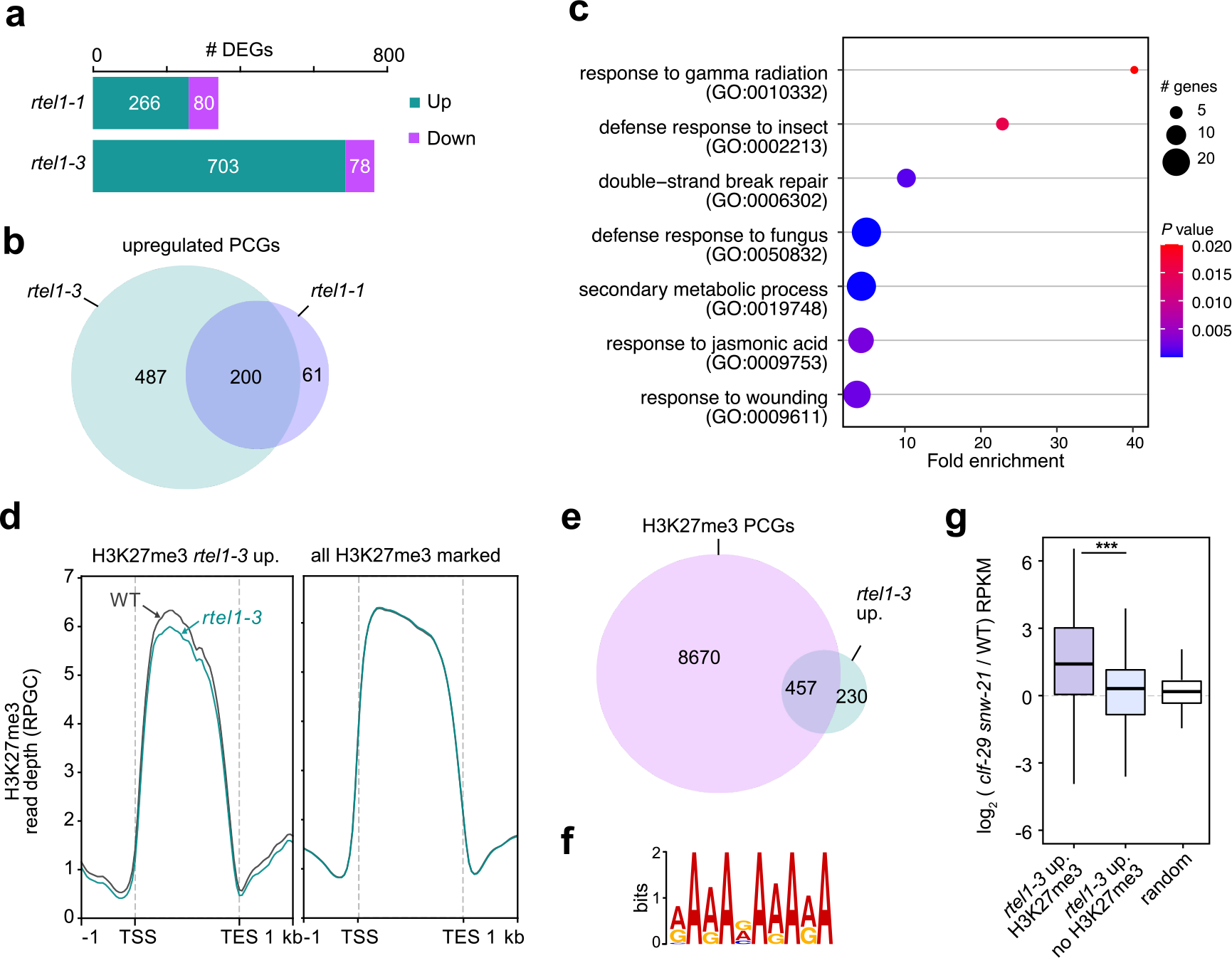
Mutations in *RTEL1* upregulate transcription of DNA damage response and H3K27me3-enriched genes. (a) Number of differentially expressed genes (DEGs) up- or down-regulated in *rtel1-1* and *rtel1-3*. **(b)** Venn diagram showing the overlap between *rte1-3* and *rtel1-1* upregulated protein-coding genes (PCGs) **(c)** Gene Ontology terms enriched in *rtel1* upregulated PCGs (common to both alleles) plotted in relation to fold enrichment. Circle size indicates the number of upregulated PCGs that are associated with each significant pathway. The circle color indicates the significant level with the adjusted *P* values < 0.05 (Fisher Exact test with Bonferroni correction, -log10). Only child GO terms are represented. **(d)** H3K27me3 ChIP signal over H3K27me3-marked PCGs upregulated in *rtel1-3* (left) and all H3K27me3-marked PCGs of the Arabidopsis genome (right). PCGs were scaled to 2 kb, and sequences 1 kb upstream their transcription start sites (TSS) or downstream their transcription end site (TES) were included. Average H3K27me3 signal over 50 bp bins is plotted. **(e)** Venn diagram showing the overlap between *rtel1-3* upregulated PCGs and all H3K27me3-marked genes. **(f)** Sequence logo of the GA-repeat motif enriched in the promoter sequences (sequences 1kb upstream of the ATG) of H3K27me3-marked *rtel1-3* upregulated PCGs. **(g)** Changes in transcript accumulation at *rtel1-3* upregulated PCGs associated or not with H3K27me3, and a random set of genes (n = 457), in *clf-29 swn-21* double mutants (raw data from (35)). Asterisks mark statistically significant differences (unpaired, two-sided Mann-Whitney test, *P* value = 1.76×10^-15^).

Previous studies in *Arabidopsis* have shown that RTEL1 is required for proper DNA replication and DNA repair (47, 48), and we found that DNA content of WT and *rtel1* mutant nuclei were similar (Supplementary Figure S3). Loss of RTEL1 causes activation of DNA repair genes (47), and accordingly, Gene Ontology (GO) analysis indicated that genes involved in response to DNA damage and DNA damage repair were overrepresented among PCGs upregulated in both *rtel1-1* and *rtel1-3* (Figure 2c, Supplementary Table S1). This suggests that *rtel1* mutants undergo constitutive replicative stress, which likely contributes to the upregulation of a subset of PCGs. Interestingly, *rtel1*-upregulated PCGs were also enriched in stress responsive genes (Figure 2c, Supplementary Table S1). Stress-responsive genes are enriched in the histone variant H2A.Z, and the majority of H2A.Z nucleosomes associate with the H3K27me3 repressive histone modification in *Arabidopsis* (49, 50). We determined genome-wide H3K27me3 patterns in wild-type and *rtel1-3* seedlings using chromatin immunoprecipitation followed by sequencing (ChIP-seq). More than 65% of *rtel1- 3* upregulated PCGs were associated with high levels of H3K27me3 in the wild type (Figure 2d, e). These PCGs H3K27me3-marked PCGs were enriched in cell cycle-regulated genes (Supplementary Figure S4) and in GO-terms related to stress response (Supplementary Tables S2, S4). Consistent with their strong association with H3K27me3, the majority (409 out of 457) of these PCGs contained at least one GA-repeat motif in their promoter sequences (Figure 2f), a motif known to act as a Polycomb response element directing the recruitement of the Polycomb repressive complex 2 (PRC2) (51). On the other hand, *rtel1-3* upregulated PCGs not marked by H3K27me3 were enriched in GO-terms involved in DNA damage response (Supplementary Tables S3, S5). The genomic distribution of *rtel1-3* differentially expressed PCGs was similar to that of all PCGs in the Arabidopsis genome, with a predominant location along chromosome arms (Supplementary Figure S5). Consistent with H3K27me3 repressing their transcription, *rtel1-3-*upregulated PCGs enriched in H3K27me3 had significantly enhanced transcript accumulation in double mutants for the CURLY LEAF and SWINGER H3K27me3 methyltransferases, as compared to *rtel1*-upregulated PCGs not marked by H3K27me3 (Figure 2g). Additionally, we found that loss of RTEL1 was associated with a moderate but reproducible reduction in H3K27me3 at upregulated PCGs associated with this mark in the WT (Figure 2d). Together, these findings suggest that both constitutive replicative stress and perturbed H3K27me3-mediated silencing likely contribute to PCG upregulation in *rtel1* through at least partly independent pathways.

### RTEL1 prevents heterochromatin DNA hypermethylation

The *QUA-QUINE STARCH* (*QQS*) gene is among the few PCGs that are downregulated in *rtel1* mutants. *QQS* is located in the pericentromeric heterochromatin of chromosome 3, and its expression is negatively correlated with the DNA methylation level of repeated sequences located within the 5’ end of the gene (52, 53). This led us to speculate that RTEL1 deficiency may alter DNA methylation patterns at some endogenous sequences. We investigated this possibility by determining genome-wide DNA methylation profiles in WT and *rtel1-3* plants using bisulfite sequencing (BS-seq). This showed that DNA methylation levels of the 5’ end of *QQS* were indeed increased in *rtel1-3* in correlation with transcriptional downregulation of the gene (Supplementary Figure S6a). More surprisingly, we found that CHG methylation levels were significantly increased at the L5-GUS transgene in *rtel1-3,* despite transcriptional reactivation (Supplementary Figure S6b).

As expected from the general anticorrelation between DNA methylation and H3K27me3 (54–56), *rtel1-3* upregulated PCGs, which are enriched in H3K27me3, showed low DNA methylation levels in the WT, and these remained unaffected in *rtel1-3* (Supplementary Figure S6c). We plotted average DNA methylation levels along chromosomes and found no overt change in CG DNA methylation in *rtel1-3* mutants (Figure 3a, Supplementary Figure S7). However, we observed that DNA methylation levels of pericentromeric regions were noticeably increased at CHG, and to a lesser extent, CHH sites (Figure 3a, Supplementary Figure S7). Consistently, average CHG, and to a lesser extent, CHH methylation levels were increased over TEs in *rtel1-3* compared to WT (Supplementary Figure S6d). All three CHG sequence subcontexts (CAG, CTG and CCG) were equally hypermethylated in *rtel1-3* (Supplementary Figure S6d). We also determined differentially methylated regions (DMRs) in *rtel1-3* relative to WT plants and found that *rtel1-3* mostly induced CHG hypermethylation, which was largely clustered at pericentromeric TE-rich heterochromatin (Figure 3c, d). Separating PCGs and TEs according to their genomic localization showed that non-CG hypermethylation in *rtel1-3* was restricted to TEs and much more pronounced at pericentromeric TEs than at TEs located on chromosome arms (Figure 3b). Analysis of the most prevalent TE superfamilies in the Arabidopsis genome, each with at least 50 copies in pericentromeres and chromosome arms, consistently showed a higher gain in CHG methylation in *rtel1-3* at pericentromeric TE copies compared to those on chromosome arms, irrespective of the TE superfamily (Supplementary Figure S8). Arabidopsis heterochromatin is associated with the H2A.W histone variant and enriched in H3K27me1 (57, 58). We found that the gain in DNA methylation in *rtel1-3* specifically occurred at regions associated with these heterochromatin features and was particularly marked at CHG sites, although heterochromatic CG and CHH sites did show some degree of hypermethylation (Figure 3e). However, this increase in DNA methylation levels was not associated with the upregulation of DNA methyltransferases genes (including *CMT3*) or the downregulation of DNA demethylase genes (Supplementary Figure S6e). These results demonstrate that RTEL1 is required for maintaining proper patterns of heterochromatin DNA methylation, particularly at CHG sites.

**Figure 3:**
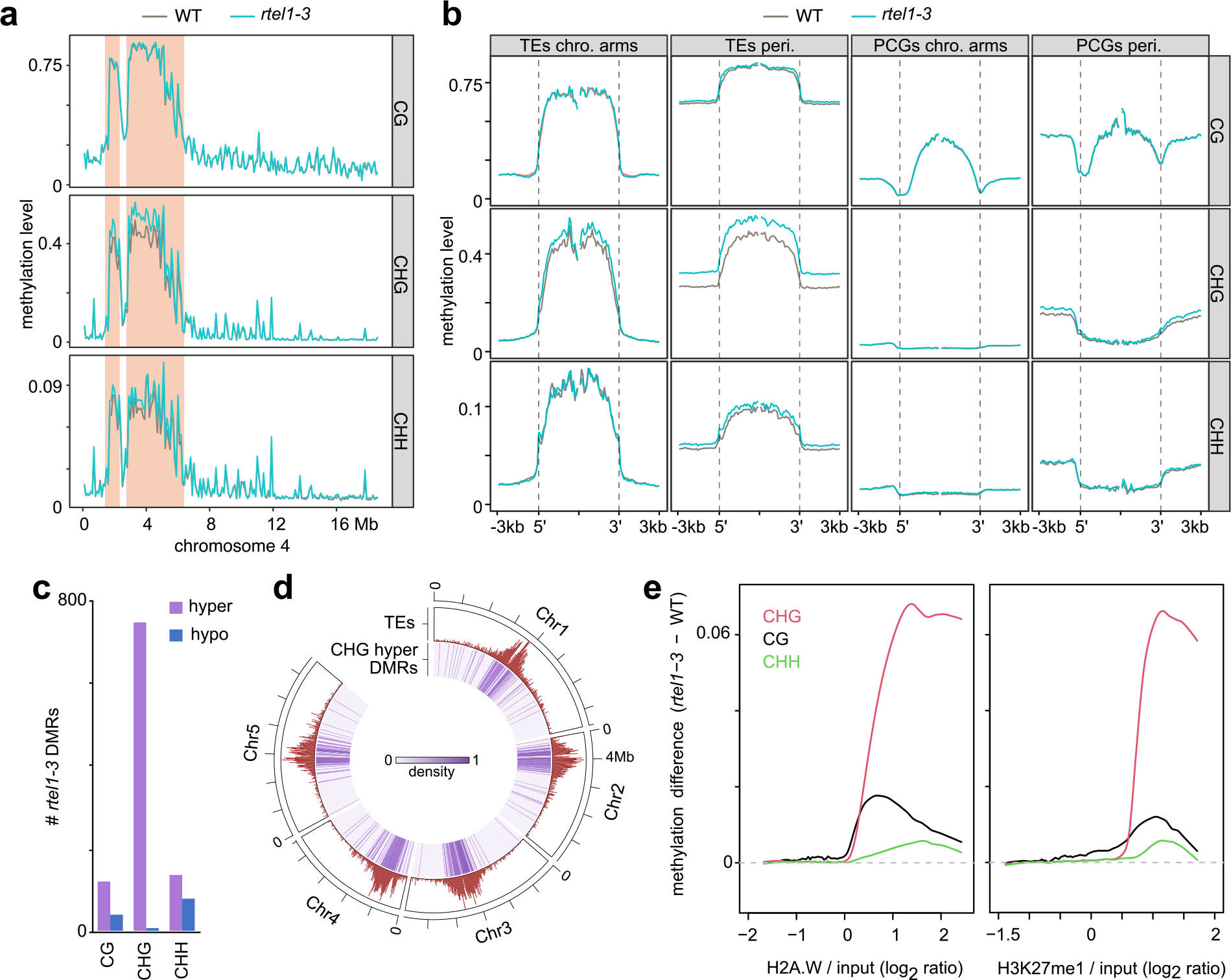
Loss of RTEL1 causes heterochromatin DNA hypermethylation. (a) Average DNA methylation level per 100kb windows on chromosome 4 in CG, CHG and CHH context. Pericentromeric heterochromatin regions are highlighted (coral color) **(b)** Metaplots showing average DNA methylation levels in all three cytosine contexts at TEs and PCGs located on chromosome arms (chro. arms) or in pericentromeric heterochromatin (peri.). TEs and PCGs were aligned at their 5’ and 3’ ends (dashed lines). Average methylation over 100bp bins 3kb upstream and 3kb downstream from the alignment points is plotted. **(c)** Number of indicated differentially methylated regions (DMRs) in *rtel1-*3. **(d)** Density of CHG hypermethylated DMRs over Arabidopsis chromosomes. Density of TEs is shown for comparison purpose. **(e)** Smoothing spline fits (50 degrees of freedom) of changes in DNA methylation (*rtel1-3* minus WT) at CG, CHG and CHH sites in 1kb windows plotted against H2A.W (left) or H3K27me1 (right) enrichment in the WT.

### Higher order chromatin organization appears unaltered in *rtel1*

In 4’,6-diamidino-2-phenylindole (DAPI) stained WT nuclei, heterochromatin appears as brightly stained foci called chromocenters, which contain most of the highly repetitive DNA in the genome, including ribosomal DNA (rDNA) and satellite repeats. The nuclei of WT and *rtel1-3* plants contained a similar number of chromocenters and an equivalent relative heterochromatic fraction (Figure 4a, b). Previous studies have shown that *rtel1-1* mutants display a reduced number of 45S rDNA repeats (48, 59), however, we were unable to detect variation in the amount of 45S rDNA repeats in *rtel1-3* (Supplementary Figure S9). To further investigate a potential impact of *rtel1-3* on higher-order chromatin structure, we profiled chromatin accessibility in WT and *rtel1-3* seedlings using Assay for Transposase-Accessible Chromatin followed by high-throughput sequencing (ATAC-seq). We found that *rtel1-3* did not affect overall chromatin accessibility profiles (Figure 4c, d). A closer examination of *rtel1* upregulated PCGs revealed that increased chromatin accessibility was not required for, nor a necessary consequence of, the transcriptional upregulation of these genes. Rather, these showed a moderate decrease in accessibility around their transcription start site (TSS), possibly reflecting active loading of the transcription machinery at these genes in the mutant (Figure 4d). In conclusion, our results suggest that higher-order chromatin organization remains largely intact in *rtel1-3*.

**Figure 4:**
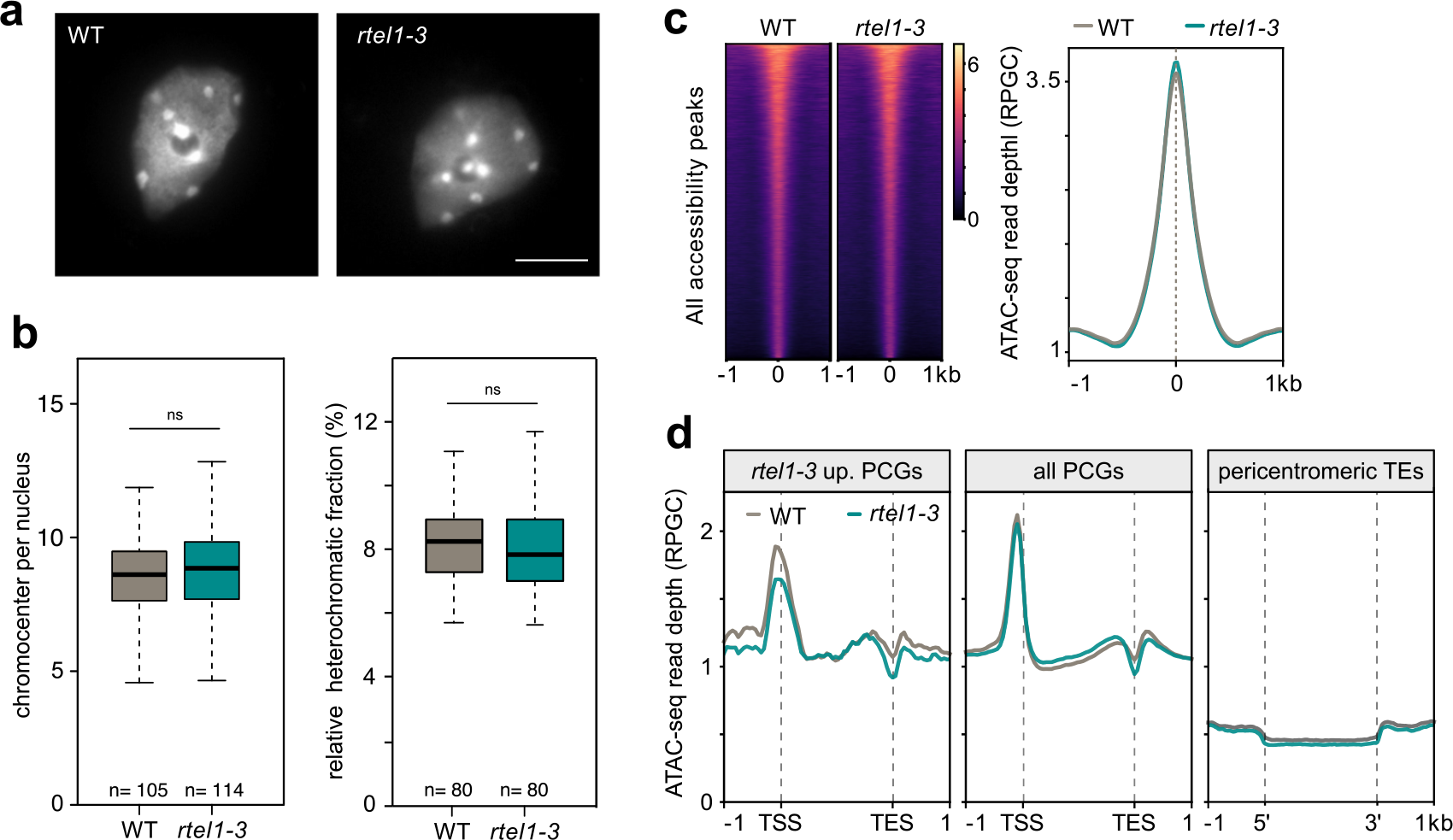
Chromatin organization appears largely unaltered in *rtel1-3*. (a) Representative photos of DAPI-stained isolated nuclei in WT (L5 background) and *rtel1-3*. Scale bar, 5µm. **(b)** Quantification of chromocenter number per nucleus (left) and heterochromatic fraction (chromocenters density / total nuclei density, right) of WT (L5 background) and *rtel1-3* plants. ns, nonsignificant (unpaired Mann-Whitney test, *P* value > 0.2). **(c)** Heatmap (left) and metaplot (right) showing ATAC-seq signal over all accessibility peaks in WT and *rtel1-3*. Peaks were aligned at their center position (0), and sequences 1kb upstream and 1kb downstream from the alignment point were included. Peaks in the heatmaps are stacked from highest (top) to lowest (bottom) ATAC-seq signal in the WT. **(d)** Metaplots showing ATAC-seq signal at *rtel1-3* upregulated PCGs, all PCGs, and pericentromeric TEs in WT and *rtel1-3*. Average signal over 100bp bins is plotted.

### RTEL1 influences gene expression largely independently of POL2A and MAIL1

Two mutant alleles of the *POL2A* gene encoding the catalytic subunit of polymerase epsilon were recovered from our screen (Supplementary Figure S1). We previously reported that *pol2a* mutations releases heterochromatin silencing and leads to increased levels of DNA methylation, particularly at CHG sites (24). However, in contrast to *rtel1-3*, TEs located outside pericentromeric heterochromatin along chromosome arms also exhibit marked CHG hypermethylation in *pol2a* mutants (Figure 3b, Supplementary Figure S10a) (24). Part of the CHG hypermethylation in *pol2a* is accounted by increased expression of *CMT3* (24); however, *CMT3* transcript accumulation was not up-regulated in *rtel1-3* (Supplementary Figure S6e). Additionally, heterochromatin gains CG methylation in *rtel1-3,* but not in *pol2a- 12* (24). We sought to analyze epistasis between *pol2a* and *rtel1* mutations. However, we were unable to isolate *rtel1 pol2a* double mutants in the progeny of self-fertilized *rtel1-3 pol2a-12* double heterozygotes (204 individuals) or either sesquimutants (100 individuals each), suggesting a lethal genetic interaction between *rtel1-3* and *pol2a-12*. Comparing the transcriptomes of *pol2a-12* and *rtel1-3* single mutants revealed little overlap of upregulated PCGs (Supplementary Figure S10b). Although perturbed H3K27me3-mediated silencing likely contributes to transcriptional upregulation of PCGs in both *pol2a* and *rtel1* (see Figure 2d-f) (24), the sets of H3K27me3-marked PCGs upregulated in each mutant were mostly distinct (Supplementary Figure S10b), and *rtel1*-upregulated PCGs displayed much higher average H3K27me3 levels than *pol2a*-upregulated genes (Supplementary Figure S10c). Altogether, our data suggest that *RTEL1* and *POL2A* function in at least partly distinct pathways to regulate gene expression and restrain DNA methylation.

We previously showed that *MAIL1* regulates the expression of numerous genes and ensures silencing of a set of pericentromeric TEs in addition to the L5-GUS transgene independently of DNA methylation (19). Although *mail1* mutants display severe developmental defects that do not resemble *rtel1-3* mutant phenotype, TEs show increased levels of non-CG methylation in both mutants (Supplementary Figure S10d) (24, 60). We sought to test the genetic interaction between *rtel1* and *mail1* mutations in non-CG DNA hypermethylation by crossing *rtel1-3* with the *mail1-1* knockout mutant. However, we were unable to obtain any *rtel1-3 mail1-1* double mutant plants in the progeny of double heterozygotes (107 individuals) or in the progeny of *rtel1-3 MAIL1/mail1-1* sesquimutant (90 individuals), indicating that RTEL1 and MAIL1 are synergistically required for viability. Transcriptome comparison of *rtel1-3* and *mail1-1* single mutants highlighted distinct sets of upregulated PCGs with limited overlap regardless of their association with H3K27me3 (Supplementary Figure S10e). We conclude that *RTEL1* and *MAIL1* regulate gene expression through largely non-overlapping molecular pathways.

### RTEL1 stabilizes H3K27me3 level over heterochromatin

We also recovered a mutant allele of *MOM1* from our screen for mutants that release silencing without loss of DNA methylation (Supplementary Figure S1). MOM1 was the first identified protein required for transcriptional silencing independent of DNA methylation in *Arabidopsis* (11), and disruption of *MOM1* releases silencing of the L5-GUS transgene and of many pericentromeric sequences (11, 13, 26). We obtained *rtel1-3 mom1-2* double mutants, which were phenotypically indistinguishable from *rtel1-3* single mutants (Figure 5a). Histochemical staining for GUS activity revealed that L5-GUS silencing was more strongly released in *mom1* than in *rtel1* and *rtel1 mom1* (Figure 5b), suggesting a genetic interaction between *rtel1* and *mom1*. Similarly, release of silencing of the *TSI* repeats was significantly stronger in *mom1-2* than in *rtel1-3,* and *rtel1 mom1* double mutants exhibited intermediate levels of *TSI* transcript accumulation (Figure 5c), indicating that *rtel1* restrains *mom1*-induced reactivation at these loci. To further investigate epistasis between *rtel1* and *mom1* mutations, we compared silencing defects in *mom1* and *mom1 rtel1* using RNA-seq. We identified 72 reactivated TEs in *mom1*, the majority of which showed a sharp reduction in transcript levels in *mom1-2 rtel1-3* (Figure 5d), suggesting that RTEL1 is required for silencing release in *mom1-2*. We previously reported that TE transcription in *mom1-2* requires MEDIATOR14 (MED14) (18). However, *MED14* transcript levels were similar in WT and *rtel1* plants (Supplementary Figure S11), suggesting that the requirement for RTEL1 and MED14 were independent. We hypothesized that RTEL1 dependency for *mom1*-induced silencing release might rely on changes in chromatin structure and/or patterns of epigenetic marks at these loci in the absence of RTEL1. Chromatin accessibility of TEs silenced by MOM1 was not altered in *rtel1* (Supplementary Figure S12). However, these TEs showed marked hypermethylation of CHG sites and, more surprisingly, an increased association with H3K27me3 in the absence of RTEL1 (Figure 5e, f). Thus, increased levels of these two repressive marks possibly contribute to limiting *mom1-*induced activation of these endogenous loci in the *rtel1 mom1* background.

**Figure 5:**
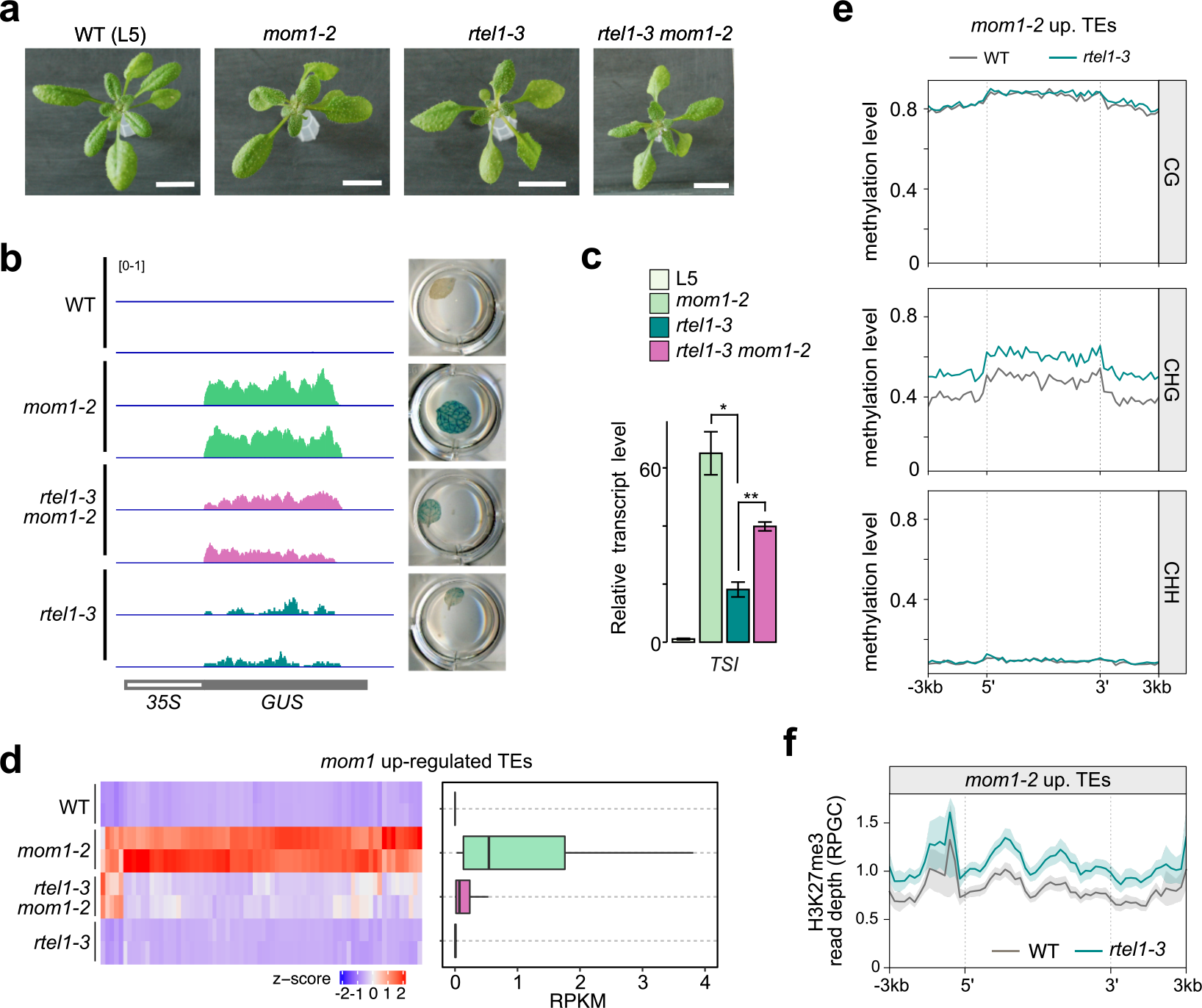
Loss of RTEL1 suppresses *mom1*-induced release of silencing. (a) Photos of 3- week-old plants of the indicated genotypes. Scale bar, 1cm. **(b)** Genome browser tracks showing mRNA profiles (RPKM) at the L5-GUS locus in the indicated genotypes (left). Two replicates are shown for mRNA-seq data. L5-GUS transgene activity detected by histochemical staining in isolated leaves of the indicated genotypes is shown on the right. **(c)** RT-qPCR analysis of *TSI* transcripts from rosette leaves of indicated genotypes. Transcript levels are normalized to *ACT2* and further normalized to L5. Sample means are shown with error bars representing standard error of the mean across three biological replicates. Asterisks mark statistically significant differences (unpaired, two-sided Student’s t-test, \**P*<0.05, \*\**P*<0.005). **(d)** Heatmap of RNA-seq expression z-scores computed for *mom1-2* upregulated TEs across the indicated genotypes (left). A boxplot with the same data expressed in RPKM is shown on the right. **(e)** Metaplots showing average DNA methylation levels in all three cytosine contexts at *mom1-2* upregulated TEs in WT and *rtel1-3*. TEs were scaled to 6 kb and sequences 3 kb upstream or downstream their 5’ and 3’ ends respectively were included. Average DNA methylation levels over 200 bp bins is plotted. **(f)** H3K27me3 ChIP-seq levels at *mom1-2* upregulated TEs in WT and *rtel1-3*. Shaded areas represent standard deviation of two biological replicates. TEs were aligned as in (e).

Since MOM1-regulated TEs are mostly located in heterochromatin, we further analyzed changes in H3K27me3 over heterochromatic sequences in *rtel1-3*. We found that, on average, TEs located in chromosome arms or in pericentromeres exhibited equally low association with H3K27me3 compared to PCGs in the WT (Figure 6a). Loss of RTEL1 was associated with increased levels of H3K27me3 over TEs and PCGs located in pericentromeres, while TEs and PCGs in chromosome arms remained largely unaffected (Figure 6a). Considering the most abundant TE superfamilies of the genome, we found that TE copies located in pericentromeres consistently showed increased H3K27me3 levels in *rtel1-3,* except for TEs of the HAT, Pogo and Helitron superfamilies (Supplementary Figure S13a). Interestingly, TEs belonging to these three superfamilies were on average short and associated with low levels of the heterochromatin marks H2A.W, H3K27me1 and H3K9me2, even when located in pericentromeric heterochromatin (Supplementary Figure S13b). PCGs located in pericentromeric regions, which gain H3K27me3 in *rtel1-3*, were also associated with higher levels of these marks compared to PCGs in chromosome arms (Figure 6a). These findings suggest that heterochromatin specifically gain H3K27me3 in *rtel1*. To strengthen this conclusion, we determined H3K27me3 changes in *rtel1* in relation to H2A.W, H3K27me1 and H3K9me2 at the whole genome level. Genomic regions depleted in these heterochromatin marks showed no change in H3K27me3, whereas enrichment in H2A.W, H3K27me1 and H3K9me2 positively correlated with with H3K27me3 gain in *rtel1* (Figure 6b). Thus, our data indicate that in the absence of RTEL1, genomic regions associated with a heterochromatin environment specifically gain H3K27me3.

**Figure 6:**
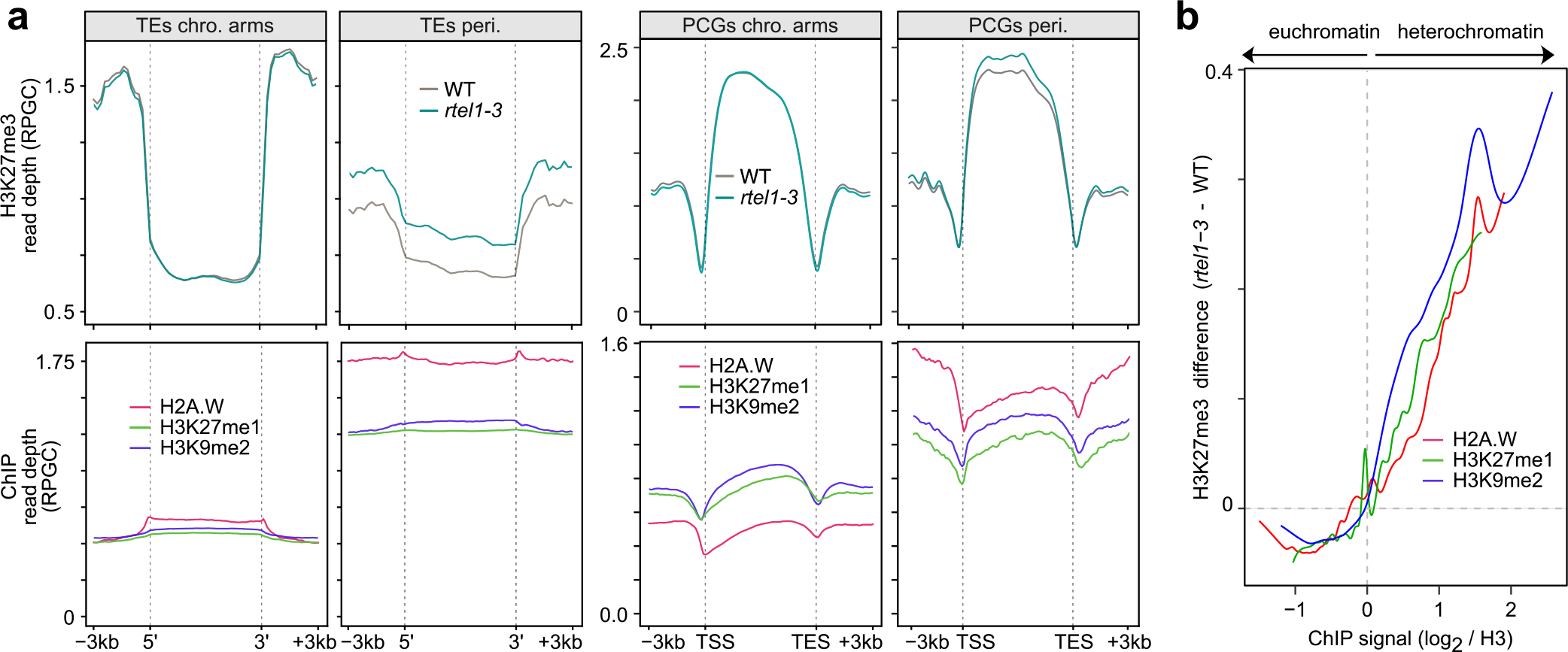
Heterochromatin H3K27me3 levels increase in *rtel1*. (a) ChIP signal for H3K27me3 in WT and *rtel1-3* (top), and ChIP signal for H2A.W, H3K27me1 and H3K9me2 in WT plants (bottom) over TEs and PCGs located in chromosome arms (chro. arms) or in pericentromeric heterochromatin (peri.). Sequences were scaled as in Figure 5e and average H3K27me3 signal over 100 bp bins is plotted. **(b)** Smoothing spline fits (50 degrees of freedom) of changes in H3K27me3 (*rtel1-3* minus WT) in 1kb windows plotted against H2A.W, H3K27me1 or H3K9me1 enrichment in the WT.

### Impact of *rtel1* at DNA replication origins

DNA replication in eukaryotes is initiated at genomic regions known as DNA replication origins. We used our BS-seq, ATAC-seq and ChIP-seq data to investigate the impact of RTEL1 loss on the chromatin environment of DNA replication origins. We found that DNA replication origins exhibited lower levels of DNA methylation in all cytosine contexts compared to flanking sequences and control (randomly shuffled) regions (Figure 7a). This observation confirms and extends a previous finding that DNA replication origins are associated with reduced CG methylation levels (61). In the *rtel1-3* mutant, both control regions and DNA replication origins gained CHG methylation; however, the magnitude of CHG hypermethylation was comparatively lower at DNA replication origins (Figure 7a). Additionally, we found that, in chromosome arms as well as in pericentromeric heterochromatin, CHG hyper-DMRs were located significantly further away from DNA replication origins than expected (Figure 7b). Therefore, we conclude that the strongest impact of RTEL1 loss on DNA methylation tends to occur outside DNA replication origins, possibly at more distant locations where replication forks progress.

**Figure 7:**
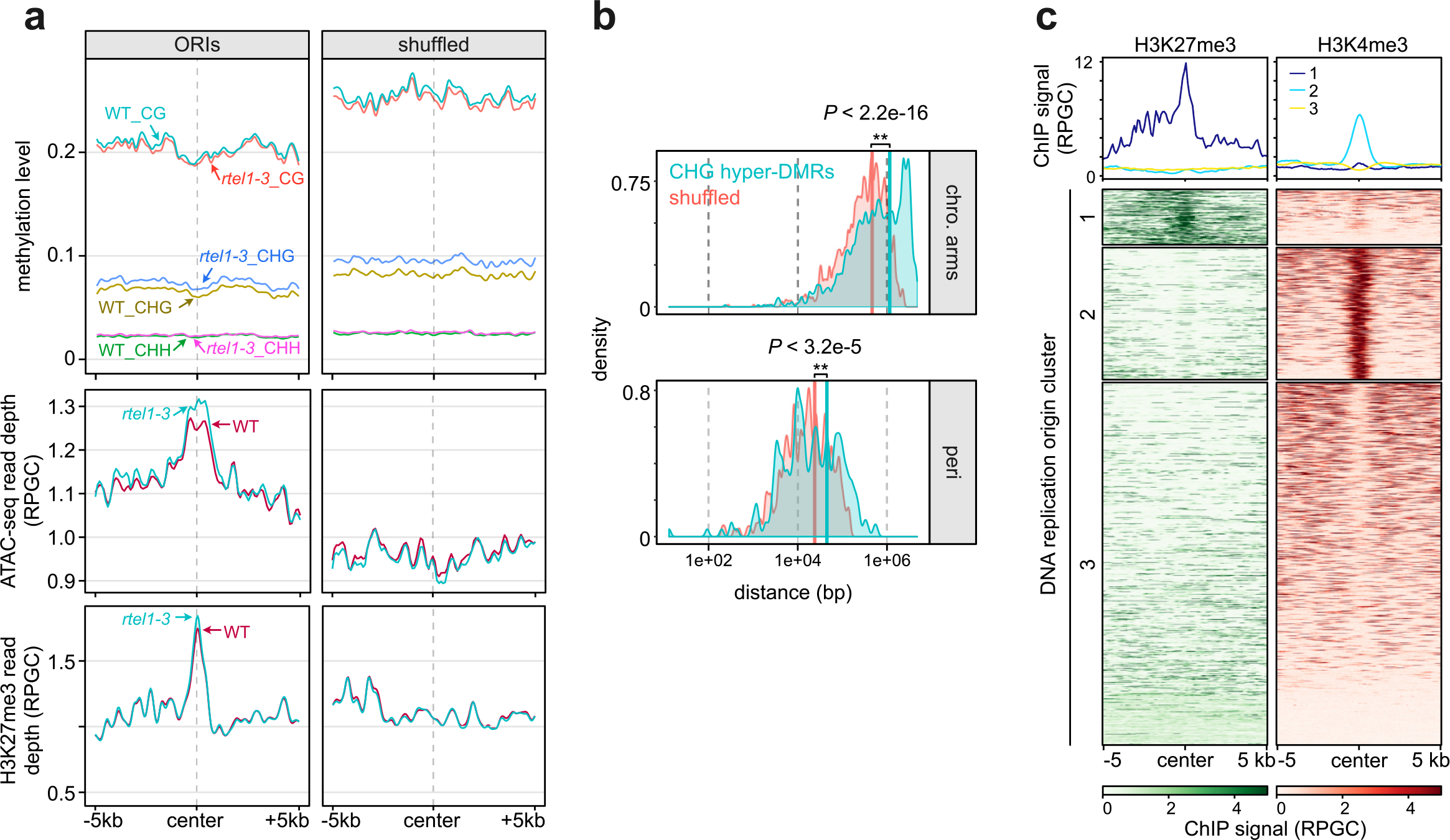
Impact of *rtel1* at DNA replication origins. (a) Metaplots showing DNA methylation, ATAC-seq signal and H3K27me3 levels at DNA replication origins (ORIs) in WT and *rtel1-3* (left). As a control, ORIs were randomly shuffled across the genome (shuffled, right). ORIs and control regions were aligned at their center position, and sequences 5kb upstream and 5kb downstream from the alignment point were included. Average signal over 100bp bins is plotted. **(b)** Density plot of the closest distance between DNA replication origins and CHG hypermethylated DMRs (light red), or randomly shuffled DMR regions (light blue). Average distances are shown as vertical bars of corresponding colors. Asterisks mark statistically significant differences (Wilcoxon signed rank test, *P* values are indicated). **(c)** K-means clustering of DNA replication origins based on H3K27me3 and H3K4me3 ChIP signals. Corresponding metaplots are shown on the top.

DNA replication origins generally exhibited greater accessibility compared to their flanking sequences and control regions, and that their accessibility slightly increased in *rtel1* (Figure 7a). We also found that DNA replication origins showed high average H3K27me3 levels (Figure 7a). This is in agreement with a previous study that showed that some DNA replication origins are associated with H3K27me3 (62). This is also consistent with the prior finding that DNA replication origins are enriched in H2A.Z, as the majority of H2A.Z nucleosomes associate with H3K27me3 (49, 57). DNA replication origins were shown to be enriched in H3K4me3 (61), and using published H3K4me3 data (63), we found that DNA replication origins marked by either H3K27me3 or by H3K4me3 predominantly form distinct clusters (Figure 7c). In the *rtel1* mutant, the level of H3K27me3 was slightly increased at DNA replication origins. Collectively, our data suggest that RTEL1 is required for maintaining proper levels of H3K27me3 and chromatin accessibility at DNA replication initiation sites. However, the most significant impact of RTEL1 loss on DNA methylation tends to occur outside DNA replication origins, possibly at more distant locations where replication forks progress.

### Accumulation of DNA cross-links destabilizes silencing

It has been shown previously that RTEL1 functions in the repair events of DNA crosslinks (47, 64). We used mitomycin C (MMC) and *cis*-platin (CP) crosslinking agents to assess whether accumulation of DNA crosslinks impacts transcriptional silencing. MMC mainly generates inter-strand cross-links on DNA, whereas CP preferentially forms intra-strand cross-links. Histochemical staining showed that applying CP or MMC to WT (L5) plants reactivated L5-GUS expression (Figure 8a, b). In these plants, L5-GUS expression appeared more pronounced in (non-replicating embryonic tissues) cotyledons, with MMC treated plants additionally showing GUS reactivation in hypocotyl and adult root regions, a pattern similar to that observed in untreated *rtel1* mutants (Figure 8a). Drug treatment did not enhance GUS reactivation in *rtel1-3* (Figure 8a, b). RNA-seq revealed that about half of the PCGs upregulated upon MMC treatment in the WT were also upregulated in untreated *rtel1-3* plants (Figure 8c). Conversely, PCGs differentially expressed in *rtel1-3* in the absence of drug were on average similarly differently expressed when WT plants were exposed to MMC (Figure 8d). Globally, transcriptional changes associated with *rtel1-3* mutation and those induced by treating WT plants with MMC showed positive correlation (Supplementary Figure S13). These data show that inducing DNA crosslinks leads to release of silencing and suggest that accumulation of these DNA lesions may be, at least partly, at the origin of transcriptional defects in *rtel1* mutants.

**Figure 8:**
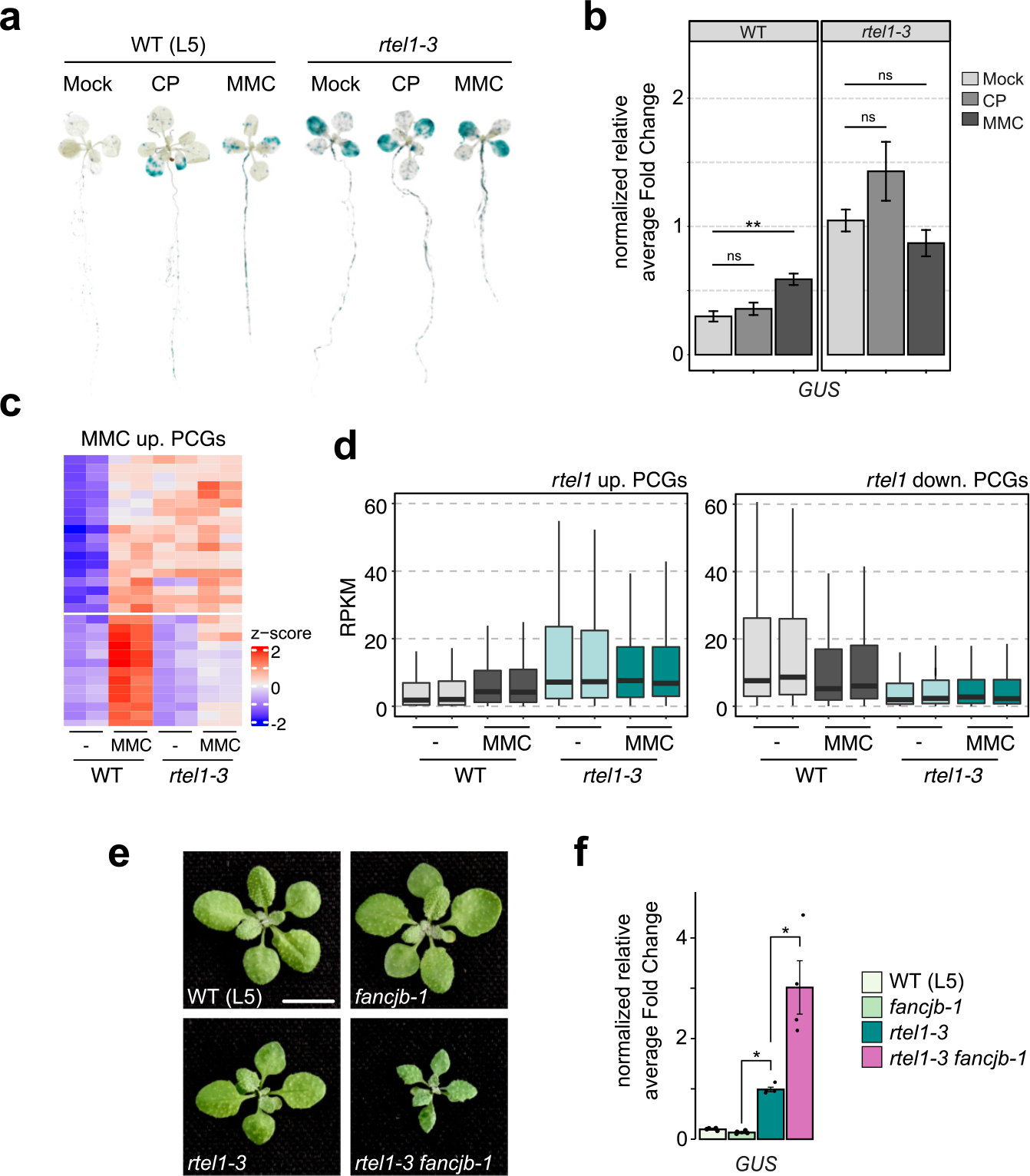
DNA cross-links accumulation destablilizes silencing. (a) L5-GUS transgene activity detected by histochemical staining in 14-day old seedlings. WT and *rtel1-3* plants were treated or not (Mock) with 2.5 mg/l of either *cis*-platin (CP) or mitomycin C (MMC) crosslinking agents. **(b)** L5-GUS transcript accumulation detected by quantitative RT-PCR in WT (L5) and *rtel1-3* seedlings untreated (mock) or treated with CP and MMC as in (a). Values represent means from two biological replicates with *rtel1-3* levels set to 1. Error bars represent standard error of the mean. Asterisks mark statistically significant differences (two- sided unpaired Mann-Whitney test, ** *P* value < 0.005); ns, nonsignificant (two-sided unpaired Mann-Whitney test, *P* value > 0.2). **(c)** Heatmap of normalized (z-score) expression level of PCGs upregulated in MMC-treated WT plants. **(d)** Box plot showing transcript accumulation (RPKM) from *rtel1* upregulated PCGs and rtel1 downregulated PCGs in WT and *rtel1-3* plants that have been treated (MMC) or not (-) with MMC. Error bars represent standard error of the mean across two biological replicates. **(e)** Representative images of 20- day old plants of the indicated genotypes. Scale bar is 1cm. **(f)** L5-GUS transcript accumulation detected by quantitative RT-PCR in the indicated genotypes. Values represent means from four biological replicates, with *rtel1-3* set to 1. Error bars represent standard error of the mean across four biological replicates. Asterisks mark statistically significant differences (two-sided unpaired Mann-Whitney test, * *P* value < 0.05).

In Arabidopsis, the FANCJB helicase, closely related to RTEL1, is also involved in the repair of inter-strand crosslinks. However, both helicases were shown to act in independent pathways of inter-strand crosslinks repair (48). We combined *rte1-3* mutation with the *fancjb- 1* mutation by crossing and obtained *rtel1-3 fancjb-1* double mutants containing the L5-GUS transgene. The *fancjb-1* mutants were indistinguishable from their WT siblings; however, *rtel1-3 fancjb-1* plants exhibited a more severe developmental phenotype compared to *rtel1- 3*and first generation double mutants were complete sterile (Figure 8e, Supplementary Figure S14a). We found that L5-GUS expression remained silenced in *fancjb-1,* whereas L5-GUS silencing was further released in *rtel1-3 fancjb-1* double mutants compared to *rtel1-3* single mutants (Figure 8f). Similarly, silencing release of the Athila retrotransposon-derivates repeats (TSI) was enhanced in *rtel1-3 fancjb-1* compared to *rtel1-3* (Supplementary Figure S14b). Thus, in the absence of RTEL1, FANCJ appears to restrain silencing release. Taken together, these results indicate that RTEL1-mediated repair of inter-strand crosslinks has a prominent role in silencing, and that FANCJB backs up RTEL1 to promote efficient silencing.

## Discussion

In this study, we uncover unappreciated roles for the DNA helicase RTEL1 in transcriptional silencing and in controlling DNA methylation and H3K27me3 patterns. RTEL1 deficiency releases transcriptional silencing and is associated with changes in H3K27me3 levels characterized by a reduction at up-regulated genes and an increase over heterochromatin. In addition, DNA methylation levels increase in heterochromatin, particularly at CHG sites.

In Arabidopsis, mouse, human cells, and *C. elegans*, RTEL1 plays an essential and conserved DNA function in facilitating replication, which ensures genome stability and cell cycle progression (47, 48, 59, 64–67). RTEL1 associates with PCNA *via* a PIP box domain in its C terminus that is conserved in human, mouse and Arabidopsis, and accumulates at sites of replication stress, suggesting that RTEL1 could target stalled replisomes and facilitate replication fork progression by resolving DNA secondary structures (64, 65). We previously reported that inhibiting replication fork progression by depleting the cellular dNTP pool through the supply of HU induces loss of silencing (24). RTEL1 deficiency is associated with prolonged S-phase duration, likely due to the accumulation of cross-linked DNA (64), and we show here that treatment with the cross-linking agents, which also likely interferes with replication fork progression, also results in release of silencing. These findings support that replication stress can trigger transcription of normally silent regions. Interestingly, we found that exposing *rtel1* plants to cross-linking agents did not further enhance the release of silencing, suggesting that *rtel1* mutation and cross-linking agents may act through a common pathway to cause silencing loss. This supports the hypothesis that RTEL1-mediated repair of inter-strand crosslinks is essential for stabilizing transcriptional silencing. FANCJB, one of the 5’-3’ DNA helicases, has been reported to play a role in repairing inter-strand crosslinks in Arabidopsis independently of RTEL1 (48). We find that the combined deficiency of both helicases results in developmental defects and complete sterility. This synthetic sterility appears to be evolutionarily conserved as *C. elegans* double mutants for *rtel1* and the FANCJ homolog *dog-1* fail to develop embryos due to germline proliferation defects, which were assigned to replicative stress (67). Our data suggest that in the absence of RTEL1, FANCJB- mediated repair of inter-strand crosslinks serves a backup function restraining loss of silencing. FANCJ was shown to function alongside RTEL1 to overcome covalent DNA-protein crosslinks that constitute bulky roadblocks to the advancing DNA replication machinery (68, 69); however, inter-strand DNA crosslinks constitute different barriers to replisome progression. Repair of covalent links between DNA strands is a complex multi-step process with several functional levels including inter-strand crosslink detection and unhooking by DNA incisions that generate a double strand break, monoadduct bypass and removing, and finally recombinational repair of double strand break, which eventually restore replication fork progression. *In vitro*, purified human RTEL1 mediates dismantling of displacement loops (D loops) that are formed as recombination intermediates (67), and this anti-recombinase activity of RTEL1, which allows replication fork progression, is conserved in Arabidopsis (47, 64, 70). Thus, RTEL1 deficiency may results in the accumulation of incompletely processed and toxic joint DNA molecules during repair of inter-strand crosslink-induced DNA damage, which perturbs DNA replication. FANCJ plays a crucial function in branch-migration, which suppresses these intermediate DNA structures (71). Additionally, FANCJ can destabilize the RAD51 nucleoprotein filament associated with single-stranded DNA and inhibit the strand exchange reaction to complete homologous recombinational repair (72). Therefore, we hypothesize that, to compensate the lack of RTEL1 and the presence of toxic recombination intermediates, FANCJB supports inter-strand crosslinks repair by inhibiting the strand exchange reaction, thereby alleviating replicative stress and silencing release.

How may replication stress destabilize silencing? Transcriptionally upregulated loci in *rtel1* are typically associated with either high levels of DNA methylation (e.g., L5-GUS transgene, TSI repeats) or enriched in H3K27me3 (PCGs). The propagation of these repressive epigenetic marks is tightly linked to DNA replication to ensure prompt restoration of the epigenetic landscape following doubling of the DNA amount (1, 73, 74). As the replication fork progresses, newly synthesized DNA strands acquire their DNA methylation pattern by copying the preexisting DNA methylation information from the template strand. Parental histones are displaced and recycled with their modifications and re-deposited at their pre-replicative positions. New, unmodified histones are deposited, resulting in a twofold dilution of parental histone modification levels. These new histones are then modified to restore the parental level of histone modifications (75, 76). In yeast and animal cells, parental histone recycling is facilitated by the tethering of parental histones to the DNA polymerases alpha and epsilon, and disruption of this interaction leads to transcriptional defects (77–79). In HeLa cells, replication stress induced by HU treatment leads to local loss of histone (H3.1) recycling and severely impairs the spatial distribution of parental histones in the surrounding regions, which may likely contribute to epigenomic instability (80). We propose that, upon RTEL1 deficiency, defective repair of inter-strand crosslink-induced DNA damage creates local DNA perturbation at the replication fork that impair recycling of parental histones and delays epigenome restoration following DNA replication. This may provide a short time-window with locally reduced levels of H3K27me3 or DNA methylation, allowing transcription factor binding and transcription initiation in typically silent genomic regions (Figure 9). The gain in H3K27me3 over heterochromatin in *rtel1* might reflect relocation of this mark from chromosome arms towards this late replicating genome compartment, which appears less affected by replication stress-induced histone recycling defects in HeLa cells (80). RTEL1 deficiency is associated with increased levels of DNA methylation in heterochromatin, most prominently at CHG sites, which occurs independently of transcriptional upregulation of the DNA methyltransferases. We previously reported that replication stress induced by HU treatment and mutations in other DNA replication-related genes, such as POL2A, RPA2A, and POLD2, also lead to increased CHG methylation levels in heterochromatin (24). Hence, heterochromatin CHG hypermethylation appears to be a molecular hallmark of replication stress. Recent studies in yeast have shown that fork stalling results in local chromatin alterations associated with higher levels of the H3K9me3 heterochromatin mark. This locally compacts chromatin and avoids fork collapse (81). In Arabidopsis, H3K9me2 serves as the functional equivalent of H3K9me3. Considering the tight interconnection between H3K9me2 and CHG methylation maintenance (82–86), a similar mechanism could contribute to the increased CHG methylation observed under replication stress. Methylation of CHG sites mostly occurs in heterochromatin, and because its maintenance is not as efficient as that of CG methylation (87), it may explain why heterochromatin hypermethylation is skewed toward CHG sites in *rtel1* and other mutants that exhibit constitutive replication stress. Heterochromatin is difficult to replicate, regularly causing replication fork stalling even in the WT (88), which could contribute to maintaining higher CHG methylation levels in these genomic regions. Increased levels of CHG methylation (and H3K27me3) in heterochromatin likely mitigate transcriptional de-repression of the underlying sequences under replication stress. This is consistent with the impact of RTEL1 deficiency on *mom1*-induced transcription of heterochromatic TEs, and with the strong upregulation of TEs we previously observed in *pol2a cmt3* double mutants (24).

**Figure 9.**
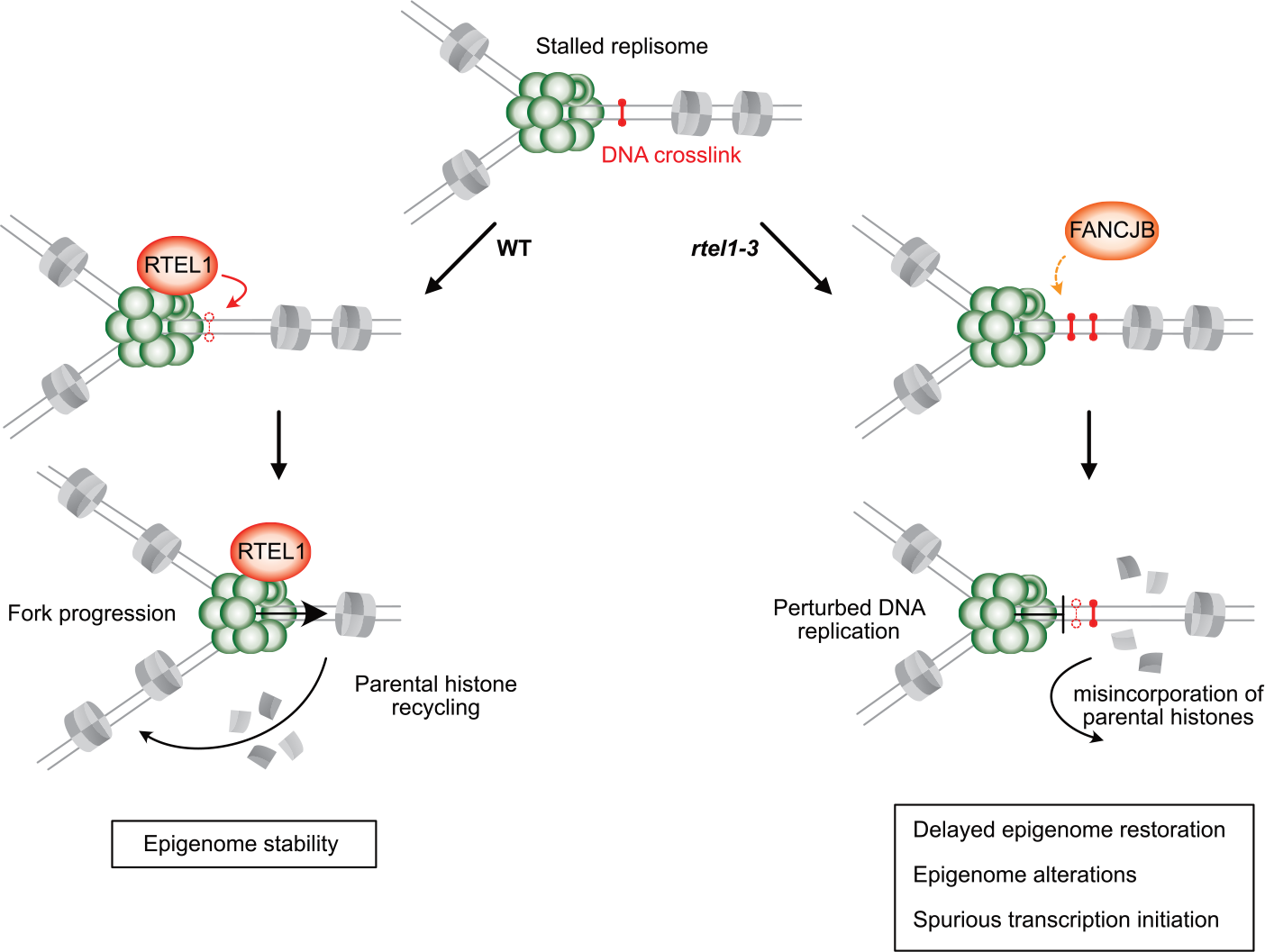
Speculative model for RTEL1 role in epigenome stability. DNA secondary structures such as DNA crosslinks can lead to replication fork stalling. The RTEL1 DNA helicase repairs these DNA lesions, allowing fork progression and proper recycling of pre-marked parental histones. In absence of RTEL1, DNA crosslink repair by FANCJB mitigates replicative stress and silencing release. However, accumulation of DNA crosslinks in absence of RTEL1 may results in perturbation of the DNA replication timing, which potentially alters recycling of parental histones and delays epigenome restoration and lead to epigenome alterations. Delayed replication fork progression may provide a temporal window for transcription factor binding and transcription initiation in typically silent genomic regions.

Our study reveals that POL2A and RTEL1 are synergistically required for Arabidopsis viability. This finding is consistent with a recent report indicating that removal of both RTEL1 and DNA polymerase epsilon leads to loss of viability in worms and vertebrates (77), indicating that this synthetic lethal interaction is conserved across different kingdoms. Notably, the combined loss of RTEL1 and DNA polymerase epsilon confers extensive genome instability and cessation of DNA replication in both worms and mouse cells (77). We also find that *rtel1* is synthetic lethal when combined with *mail1*, which is known to cause DNA damage accumulation (89). This suggests that RTEL1 function is essential to DNA replication in the presence of DNA damage. Together, our results highlight the importance of RTEL1 and its interaction with other DNA replication factors in maintaining epigenome integrity and cell viability.

## Data Availability

High throughput sequencing data generated in this study has been deposited in the Gene Expression Omnibus (GEO) database under accession number GSE197614. All other data supporting the findings of this study are available within the manuscript and its Supplementary Files or are available from the corresponding author upon request.

## Acknowledgments

We thank Hervé Vaucheret for the kind gift of the L5 line.

## References

1. Law, J.A. and Jacobsen, S.E. (2010) Establishing, maintaining and modifying DNA methylation patterns in plants and animals. Nat. Rev. Genet., 11, 204–220.

2. Tran, R.K., Henikoff, J.G., Zilberman, D., Ditt, R.F., Jacobsen, S.E. and Henikoff, S. (2005) DNA Methylation Profiling Identifies CG Methylation Clusters in Arabidopsis Genes. Curr. Biol., 15, 154–159.

3. Zhang, X., Yazaki, J., Sundaresan, A., Cokus, S., Chan, S.W.-L., Chen, H., Henderson, I.R., Shinn, P., Pellegrini, M., Jacobsen, S.E., et al. (2006) Genome-wide High-Resolution Mapping and Functional Analysis of DNA Methylation in Arabidopsis. Cell, 126, 1189–1201.

4. Lister, R., O’Malley, R.C., Tonti-Filippini, J., Gregory, B.D., Berry, C.C., Millar, A.H. and Ecker, J.R. (2008) Highly Integrated Single-Base Resolution Maps of the Epigenome in Arabidopsis. Cell, 133, 523–536.

5. Bieluszewski, T., Xiao, J., Yang, Y. and Wagner, D. (2021) PRC2 activity, recruitment, and silencing: a comparative perspective. Trends Plant Sci., 26, 1186–1198.

6. Xiao, J. and Wagner, D. (2015) Polycomb repression in the regulation of growth and development in Arabidopsis. Curr. Opin. Plant Biol., 23, 15–24.

7. Du, J., Johnson, L.M., Jacobsen, S.E. and Patel, D.J. (2015) DNA methylation pathways and their crosstalk with histone methylation. Nat. Rev. Mol. Cell Biol., 16, 519–532.

8. Zemach, A., Kim, M.Y., Hsieh, P.-H., Coleman-Derr, D., Eshed-Williams, L., Thao, K., Harmer, S.L. and Zilberman, D. (2013) The nucleosome remodeler DDM1 allows DNA methyltransferases to access H1-containing heterochromatin. Cell, 153, 193–205.

9. Vongs, A., Kakutani, T., Martienssen, R.A. and Richards, E.J. (1993) Arabidopsis thaliana DNA Methylation Mutants. Science, 260, 1926–1928.

10. Mathieu, O., Reinders, J., Čaikovski, M., Smathajitt, C. and Paszkowski, J. (2007) Transgenerational Stability of the Arabidopsis Epigenome Is Coordinated by CG Methylation. Cell, 130, 851–862.

11. Amedeo, P., Habu, Y., Afsar, K., Scheid, O.M. and Paszkowski, J. (2000) Disruption of the plant gene MOM releases transcriptional silencing of methylated genes. Nature, 405, 203– 206.

12. Steimer, A., Amedeo, P., Afsar, K., Fransz, P., Scheid, O.M. and Paszkowski, J. (2000) Endogenous Targets of Transcriptional Gene Silencing in Arabidopsis. Plant Cell, 12, 1165– 1178.

13. Vaillant, I., Schubert, I., Tourmente, S. and Mathieu, O. (2006) MOM1 mediates DNA-methylation-independent silencing of repetitive sequences in Arabidopsis. EMBO Rep., 7, 1273–1278.

14. Yokthongwattana, C., Bucher, E., Čaikovski, M., Vaillant, I., Nicolet, J., Scheid, O.M. and Paszkowski, J. (2010) MOM1 and Pol-IV/V interactions regulate the intensity and specificity of transcriptional gene silencing. EMBO J., 29, 340–351.

15. Moissiard, G., Cokus, S.J., Cary, J., Feng, S., Billi, A.C., Stroud, H., Husmann, D., Zhan, Y., Lajoie, B.R., McCord, R.P., et al. (2012) MORC Family ATPases Required for Heterochromatin Condensation and Gene Silencing. Science, 336, 1448–1451.

16. Moissiard, G., Bischof, S., Husmann, D., Pastor, W.A., Hale, C.J., Yen, L., Stroud, H., Papikian, A., Vashisht, A.A., Wohlschlegel, J.A., et al. (2014) Transcriptional gene silencing by Arabidopsis microrchidia homologues involves the formation of heteromers. Proc. Natl. Acad. Sci., 111, 7474–7479.

17. Li, Z., Wang, M., Zhong, Z., Gallego-Bartolomé,J., Feng, S., Jami-Alahmadi, Y., Wang, X., Wohlschlegel, J., Bischof, S., Long, J.A., et al. (2023) The MOM1 complex recruits the RdDM machinery via MORC6 to establish de novo DNA methylation. 10.1101/2023.01.10.523455.

18. Bourguet, P., de Bossoreille, S., López-González, L., Pouch-Pélissier, M.-N., Gómez-Zambrano,Á., Devert, A., Pélissier, T., Pogorelcnik, R., Vaillant, I. and Mathieu, O. (2018) A role for MED14 and UVH6 in heterochromatin transcription upon destabilization of silencing. Life Sci. Alliance, 1, e201800197.

19. Ikeda, Y., Pélissier, T., Bourguet, P., Becker, C., Pouch-Pélissier, M.-N., Pogorelcnik, R., Weingartner, M., Weigel, D., Deragon, J.-M. and Mathieu, O. (2017) Arabidopsis proteins with a transposon-related domain act in gene silencing. Nat. Commun., 8, 15122.

20. Probst, A.V., Dunleavy, E. and Almouzni, G. (2009) Epigenetic inheritance during the cell cycle. Nat. Rev. Mol. Cell Biol., 10, 192–206.

21. Sarkies, P. and Sale, J.E. (2012) Propagation of histone marks and epigenetic memory during normal and interrupted DNA replication. Cell. Mol. Life Sci., 69, 697–716.

22. Liu, J., Ren, X., Yin, H., Wang, Y., Xia, R., Wang, Y. and Gong, Z. (2010) Mutation in the catalytic subunit of DNA polymerase α influences transcriptional gene silencing and homologous recombination in Arabidopsis. Plant J., 61, 36–45.

23. Iglesias, F.M., Bruera, N.A., Dergan-Dylon, S., Marino-Buslje, C., Lorenzi, H., Mateos, J.L., Turck, F., Coupland, G. and Cerdán, P.D. (2015) The Arabidopsis DNA Polymerase δ Has a Role in the Deposition of Transcriptionally Active Epigenetic Marks, Development and Flowering. PLoS Genet., 11, e1004975.

24. Bourguet, P., López-González, L., Gómez-Zambrano,Á., Pélissier, T., Hesketh, A., Potok, M.E., Pouch-Pélissier, M.-N., Perez, M., Da Ines, O., Latrasse, D., et al. (2020) DNA polymerase epsilon is required for heterochromatin maintenance in Arabidopsis. Genome Biol., 21, 283.

25. Yin, H., Zhang, X., Liu, J., Wang, Y., He, J., Yang, T., Hong, X., Yang, Q. and Gong, Z. (2009) Epigenetic Regulation, Somatic Homologous Recombination, and Abscisic Acid Signaling Are Influenced by DNA Polymerase ε Mutation in Arabidopsis. Plant Cell, 21, 386–402.

26. Elmayan, T., Proux, F. and Vaucheret, H. (2005) Arabidopsis RPA2: A Genetic Link among Transcriptional Gene Silencing, DNA Repair, and DNA Replication. Curr. Biol., 15, 1919– 1925.

27. Takeda, S., Tadele, Z., Hofmann, I., Probst, A.V., Angelis, K.J., Kaya, H., Araki, T., Mengiste, T., Scheid, O.M., Shibahara, K., et al. (2004) BRU1, a novel link between responses to DNA damage and epigenetic gene silencing in Arabidopsis. Genes Dev., 18, 782–793.

28. Zhang, J., Xie, S., Zhu, J.-K. and Gong, Z. (2016) Requirement for flap endonuclease 1 (FEN1) to maintain genomic stability and transcriptional gene silencing in Arabidopsis. *Plant J*. Cell Mol. Biol., 87, 629–640.

29. Schönrock, N., Exner, V., Probst, A., Gruissem, W. and Hennig, L. (2006) Functional Genomic Analysis of CAF-1 Mutants in Arabidopsis thaliana. J. Biol. Chem., 281, 9560– 9568.

30. Li, H. and Durbin, R. (2009) Fast and accurate short read alignment with Burrows–Wheeler transform. Bioinformatics, 25, 1754–1760.

31. Livak, K.J. and Schmittgen, T.D. (2001) Analysis of Relative Gene Expression Data Using Real-Time Quantitative PCR and the 2−ΔΔCT Method. Methods, 25, 402–408.

32. Dobin, A., Davis, C.A., Schlesinger, F., Drenkow, J., Zaleski, C., Jha, S., Batut, P., Chaisson, M. and Gingeras, T.R. (2013) STAR: ultrafast universal RNA-seq aligner. Bioinformatics, 29, 15–21.

33. Liao, Y., Smyth, G.K. and Shi, W. (2014) featureCounts: an efficient general purpose program for assigning sequence reads to genomic features. Bioinformatics, 30, 923–930.

34. Love, M.I., Huber, W. and Anders, S. (2014) Moderated estimation of fold change and dispersion for RNA-seq data with DESeq2. Genome Biol., 15, 550.

35. Wang, H., Liu, C., Cheng, J., Liu, J., Zhang, L., He, C., Shen, W.-H., Jin, H., Xu, L. and Zhang, Y. (2016) Arabidopsis Flower and Embryo Developmental Genes are Repressed in Seedlings by Different Combinations of Polycomb Group Proteins in Association with Distinct Sets of Cis-regulatory Elements. PLoS Genet., 12, e1005771.

36. Mi, H., Muruganujan, A., Ebert, D., Huang, X. and Thomas, P.D. (2019) PANTHER version 14: more genomes, a new PANTHER GO-slim and improvements in enrichment analysis tools. Nucleic Acids Res., 47, D419–D426.

37. Ashburner, M., Ball, C.A., Blake, J.A., Botstein, D., Butler, H., Cherry, J.M., Davis, A.P., Dolinski, K., Dwight, S.S., Eppig, J.T., et al. (2000) Gene Ontology: tool for the unification of biology. Nat. Genet., 25, 25–29.

38. The Gene Ontology Consortium, Carbon, S., Douglass, E., Good, B.M., Unni, D.R., Harris, N.L., Mungall, C.J., Basu, S., Chisholm, R.L., Dodson, R.J., et al. (2021) The Gene Ontology resource: enriching a GOld mine. Nucleic Acids Res., 49, D325–D334.

39. Schultz, M.D., He, Y., Whitaker, J.W., Hariharan, M., Mukamel, E.A., Leung, D., Rajagopal, N., Nery, J.R., Urich, M.A., Chen, H., et al. (2015) Human Body Epigenome Maps Reveal Noncanonical DNA Methylation Variation. Nature, 523, 212–216.

40. Langmead, B. and Salzberg, S.L. (2012) Fast gapped-read alignment with Bowtie 2. Nat. Methods, 9, 357–359.

41. Bourguet, P., Picard, C.L., Yelagandula, R., Pélissier, T., Lorković,Z.J., Feng, S., Pouch-Pélissier, M.-N., Schmücker, A., Jacobsen, S.E., Berger, F., et al. (2021) The histone variant H2A.W and linker histone H1 co-regulate heterochromatin accessibility and DNA methylation. Nat. Commun., 12, 2683.

42. Yelagandula, R., Osakabe, A., Axelsson, E., Berger, F. and Kawashima, T. (2017) Genome-Wide Profiling of Histone Modifications and Histone Variants in Arabidopsis thaliana and Marchantia polymorpha. In Busch, W. (ed), Plant Genomics: Methods and Protocols, Methods in Molecular Biology. Springer, New York, NY, pp. 93–106.

43. Villar, C.B.R. and Köhler, C. (2010) Plant Chromatin Immunoprecipitation. In Hennig, L., Köhler, C. (eds), Plant Developmental Biology: Methods and Protocols, Methods in Molecular Biology. Humana Press, Totowa, NJ, pp. 401–411.

44. Zhang, Y., Liu, T., Meyer, C.A., Eeckhoute, J., Johnson, D.S., Bernstein, B.E., Nusbaum, C., Myers, R.M., Brown, M., Li, W., et al. (2008) Model-based Analysis of ChIP-Seq (MACS). Genome Biol., 9, R137.

45. Ramírez, F., Ryan, D.P., Grüning, B., Bhardwaj, V., Kilpert, F., Richter, A.S., Heyne, S., Dündar, F. and Manke, T. (2016) deepTools2: a next generation web server for deep-sequencing data analysis. Nucleic Acids Res., 44, W160–W165.

46. Morel, J.-B., Mourrain, P., Béclin, C. and Vaucheret, H. (2000) DNA methylation and chromatin structure affect transcriptional and post-transcriptional transgene silencing in Arabidopsis. Curr. Biol., 10, 1591–1594.

47. Recker, J., Knoll, A. and Puchta, H. (2014) The Arabidopsis thaliana Homolog of the Helicase RTEL1 Plays Multiple Roles in Preserving Genome Stability. Plant Cell, 26, 4889– 4902.

48. Dorn, A., Feller, L., Castri, D., Röhrig, S., Enderle, J., Herrmann, N.J., Block-Schmidt, A., Trapp, O., Köhler, L. and Puchta, H. (2019) An Arabidopsis FANCJ helicase homologue is required for DNA crosslink repair and rDNA repeat stability. PLOS Genet., 15, e1008174.

49. Carter, B., Bishop, B., Ho, K.K., Huang, R., Jia, W., Zhang, H., Pascuzzi, P.E., Deal, R.B. and Ogas, J. (2018) The Chromatin Remodelers PKL and PIE1 Act in an Epigenetic Pathway That Determines H3K27me3 Homeostasis in Arabidopsis. Plant Cell, 30, 1337–1352.

50. Coleman-Derr, D. and Zilberman, D. (2012) Deposition of Histone Variant H2A.Z within Gene Bodies Regulates Responsive Genes. PLoS Genet., 8, e1002988.

51. Xiao, J., Jin, R., Yu, X., Shen, M., Wagner, J.D., Pai, A., Song, C., Zhuang, M., Klasfeld, S., He, C., et al. (2017) Cis and trans determinants of epigenetic silencing by Polycomb repressive complex 2 in Arabidopsis. Nat. Genet. 2017 4910, 49, 1546–1552.

52. Silveira, A.B., Trontin, C., Cortijo, S., Barau, J., Del Bem, L.E.V., Loudet, O., Colot, V. and Vincentz, M. (2013) Extensive Natural Epigenetic Variation at a De Novo Originated Gene. PLoS Genet., 9, e1003437.

53. Rigal, M., Becker, C., Pélissier, T., Pogorelcnik, R., Devos, J., Ikeda, Y., Weigel, D. and Mathieu, O. (2016) Epigenome confrontation triggers immediate reprogramming of DNA methylation and transposon silencing in Arabidopsis thaliana F1 epihybrids. Proc. Natl. Acad. Sci. U. S. A., 113, E2083–E2092.

54. Mathieu, O., Probst, A.V. and Paszkowski, J. (2005) Distinct regulation of histone H3 methylation at lysines 27 and 9 by CpG methylation in Arabidopsis. EMBO J., 24, 2783– 2791.

55. Deleris, A., Stroud, H., Bernatavichute, Y., Johnson, E., Klein, G., Schubert, D. and Jacobsen, S.E. (2012) Loss of the DNA Methyltransferase MET1 Induces H3K9 Hypermethylation at PcG Target Genes and Redistribution of H3K27 Trimethylation to Transposons in Arabidopsis thaliana. PLoS Genet., 8, e1003062.

56. Weinhofer, I., Hehenberger, E., Roszak, P., Hennig, L. and Köhler, C. (2010) H3K27me3 Profiling of the Endosperm Implies Exclusion of Polycomb Group Protein Targeting by DNA Methylation. PLoS Genet., 6, e1001152.

57. Yelagandula, R., Stroud, H., Holec, S., Zhou, K., Feng, S., Zhong, X., Muthurajan, U.M., Nie, X., Kawashima, T., Groth, M., et al. (2014) The Histone Variant H2A.W Defines Heterochromatin and Promotes Chromatin Condensation in Arabidopsis. Cell, 158, 98–109.

58. Jacob, Y., Feng, S., LeBlanc, C.A., Bernatavichute, Y.V., Stroud, H., Cokus, S., Johnson, L.M., Pellegrini, M., Jacobsen, S.E. and Michaels, S.D. (2009) ATXR5 and ATXR6 are novel H3K27 monomethyltransferases required for chromatin structure and gene silencing. Nat. Struct. Mol. Biol., 16, 763–768.

59. Röhrig, S., Schröpfer, S., Knoll, A. and Puchta, H. (2016) The RTR Complex Partner RMI2 and the DNA Helicase RTEL1 Are Both Independently Involved in Preserving the Stability of 45S rDNA Repeats in Arabidopsis thaliana. PLoS Genet., 12, e1006394.

60. Nicolau, M., Picault, N., Descombin, J., Jami-Alahmadi, Y., Feng, S., Bucher, E., Jacobsen, S.E., Deragon, J.-M., Wohlschlegel, J. and Moissiard, G. (2020) The plant mobile domain proteins MAIN and MAIL1 interact with the phosphatase PP7L to regulate gene expression and silence transposable elements in Arabidopsis thaliana. PLoS Genet., 16, e1008324.

61. Costas, C., de la Paz Sanchez, M., Stroud, H., Yu, Y., Oliveros, J.C., Feng, S., Benguria, A., López-Vidriero, I., Zhang, X., Solano, R., et al. (2011) Genome-wide mapping of Arabidopsis origins of DNA replication and their associated epigenetic marks. Nat. Struct. Mol. Biol., 18, 395–400.

62. Sequeira-Mendes, J., Vergara, Z., Peiró,R., Morata, J., Aragüez, I., Costas, C., Mendez-Giraldez, R., Casacuberta, J.M., Bastolla, U. and Gutierrez, C. (2019) Differences in firing efficiency, chromatin, and transcription underlie the developmental plasticity of the Arabidopsis DNA replication origins. Genome Res., 29, 784–797.

63. Liu, Q., Bischof, S., Harris, C.J., Zhong, Z., Zhan, L., Nguyen, C., Rashoff, A., Barshop, W.D., Sun, F., Feng, S., et al. (2020) The characterization of Mediator 12 and 13 as conditional positive gene regulators in Arabidopsis. Nat. Commun., 11, 2798.

64. Hu, Z., Cools, T., Kalhorzadeh, P., Heyman, J. and De Veylder, L. (2015) Deficiency of the Arabidopsis Helicase RTEL1 Triggers a SOG1-Dependent Replication Checkpoint in Response to DNA Cross-Links. Plant Cell, 27, 149–161.

65. Vannier, J.-B., Sandhu, S., Petalcorin, M.IR., Wu, X., Nabi, Z., Ding, H. and Boulton, S.J. (2013) RTEL1 Is a Replisome-Associated Helicase That Promotes Telomere and Genome-Wide Replication. Science, 342, 239–242.

66. Uringa, E.-J., Lisaingo, K., Pickett, H.A., Brind’Amour, J., Rohde, J.-H., Zelensky, A., Essers, J. and Lansdorp, P.M. (2012) RTEL1 contributes to DNA replication and repair and telomere maintenance. Mol. Biol. Cell, 23, 2782–2792.

67. Barber, L.J., Youds, J.L., Ward, J.D., McIlwraith, M.J., O’Neil, N.J., Petalcorin, M.I.R., Martin, J.S., Collis, S.J., Cantor, S.B., Auclair, M., et al. (2008) RTEL1 Maintains Genomic Stability by Suppressing Homologous Recombination. Cell, 135, 261–271.

68. Sparks, J.L., Chistol, G., Gao, A.O., Räschle, M., Larsen, N.B., Mann, M., Duxin, J.P. and Walter, J.C. (2019) The CMG Helicase Bypasses DNA-Protein Cross-Links to Facilitate Their Repair. Cell, 176, 167–181.e21.

69. Yaneva, D., Sparks, J.L., Donsbach, M., Zhao, S., Weickert, P., Bezalel-Buch, R., Stingele, J. and Walter, J.C. (2023) The FANCJ helicase unfolds DNA-protein crosslinks to promote their repair. Mol. Cell, 83, 43–56.e10.

70. Olivier, M., Charbonnel, C., Amiard, S., White, C.I. and Gallego, M.E. (2018) RAD51 and RL1 compensate telomere loss in the absence of telomerase. Nucleic Acids Res., 46, 2432.

71. Awate, S., Sommers, J.A., Datta, A., Nayak, S., Bellani, M.A., Yang, O., Dunn, C.A., Nicolae, C.M., Moldovan, G.-L., Seidman, M.M., et al. (2020) FANCJ compensates for RAP80 deficiency and suppresses genomic instability induced by interstrand cross-links. Nucleic Acids Res., 48, 9161–9180.

72. Sommers, J.A., Rawtani, N., Gupta, R., Bugreev, D.V., Mazin, A.V., Cantor, S.B. and Brosh, R.M. (2009) FANCJ Uses Its Motor ATPase to Destabilize Protein-DNA Complexes, Unwind Triplexes, and Inhibit RAD51 Strand Exchange. J. Biol. Chem., 284, 7505–7517.

73. Jiang, D. and Berger, F. (2017) DNA replication–coupled histone modification maintains Polycomb gene silencing in plants. Science, 357, 1146–1149.

74. Xu, M., Long, C., Chen, X., Huang, C., Chen, S. and Zhu, B. (2010) Partitioning of Histone H3-H4 Tetramers During DNA Replication–Dependent Chromatin Assembly. Science, 328, 94–98.

75. Alabert, C., Barth, T.K., Reverón-Gómez, N., Sidoli, S., Schmidt, A., Jensen, O.N., Imhof, A. and Groth, A. (2015) Two distinct modes for propagation of histone PTMs across the cell cycle. Genes Dev., 29, 585–590.

76. Reverón-Gómez, N., González-Aguilera, C., Stewart-Morgan, K.R., Petryk, N., Flury, V., Graziano, S., Johansen, J.V., Jakobsen, J.S., Alabert, C. and Groth, A. (2018) Accurate Recycling of Parental Histones Reproduces the Histone Modification Landscape during DNA Replication. Mol. Cell, 72, 239–249.e5.

77. Bellelli, R., Youds, J., Borel, V., Svendsen, J., Pavicic-Kaltenbrunner, V. and Boulton, S.J. (2020) Synthetic Lethality between DNA Polymerase Epsilon and RTEL1 in Metazoan DNA Replication. Cell Rep., 31, 107675.

78. Evrin, C., Maman, J.D., Diamante, A., Pellegrini, L. and Labib, K. (2018) Histone H2ALH2B binding by Pol α in the eukaryotic replisome contributes to the maintenance of repressive chromatin. EMBO J., 37, e99021.

79. Li, Z., Hua, X., Serra-Cardona, A., Xu, X., Gan, S., Zhou, H., Yang, W.-S., Chen, C., Xu, R.-M. and Zhang, Z. (2020) DNA polymerase α interacts with H3-H4 and facilitates the transfer of parental histones to lagging strands. Sci. Adv., 6, eabb5820.

80. Clément, C., Orsi, G.A., Gatto, A., Boyarchuk, E., Forest, A., Hajj, B., Miné-Hattab, J., Garnier, M., Gurard-Levin, Z.A., Quivy, J.-P., et al. (2018) High-resolution visualization of H3 variants during replication reveals their controlled recycling. Nat. Commun., 9, 3181.

81. Feng, G., Yuan, Y., Li, Z., Wang, L., Zhang, B., Luo, J., Ji, J. and Kong, D. (2019) Replication fork stalling elicits chromatin compaction for the stability of stalling replication forks. Proc. Natl. Acad. Sci., 116, 14563–14572.

82. Stroud, H., Do, T., Du, J., Zhong, X., Feng, S., Johnson, L., Patel, D.J. and Jacobsen, S.E. (2014) Non-CG methylation patterns shape the epigenetic landscape in Arabidopsis. Nat. Struct. Mol. Biol., 21, 64–72.

83. Du, J., Zhong, X., Bernatavichute, Y.V., Stroud, H., Feng, S., Caro, E., Vashisht, A.A., Terragni, J., Chin, H.G., Tu, A., et al. (2012) Dual Binding of Chromomethylase Domains to H3K9me2-Containing Nucleosomes Directs DNA Methylation in Plants. Cell, 151, 167–180.

84. Du, J., Johnson, L.M., Groth, M., Feng, S., Hale, C.J., Li, S., Vashisht, A.A., Gallego-Bartolome, J., Wohlschlegel, J.A., Patel, D.J., et al. (2014) Mechanism of DNA methylation-directed histone methylation by KRYPTONITE. Mol. Cell, 55, 495–504.

85. Johnson, L.M., Bostick, M., Zhang, X., Kraft, E., Henderson, I., Callis, J. and Jacobsen, S.E. (2007) The SRA Methyl-Cytosine-Binding Domain Links DNA and Histone Methylation. Curr. Biol., 17, 379–384.

86. Li, X., Harris, C.J., Zhong, Z., Chen, W., Liu, R., Jia, B., Wang, Z., Li, S., Jacobsen, S.E. and Du, J. (2018) Mechanistic insights into plant SUVH family H3K9 methyltransferases and their binding to context-biased non-CG DNA methylation. Proc. Natl. Acad. Sci., 115, E8793– E8802.

87. Niederhuth, C.E., Bewick, A.J., Ji, L., Alabady, M.S., Kim, K.D., Li, Q., Rohr, N.A., Rambani, A., Burke, J.M., Udall, J.A., et al. (2016) Widespread natural variation of DNA methylation within angiosperms. Genome Biol., 17, 194.

88. Zeman, M.K. and Cimprich, K.A. (2014) Causes and Consequences of Replication Stress. Nat. Cell Biol., 16, 2–9.

89. de Luxán-Hernández, C., Lohmann, J., Hellmeyer, W., Seanpong, S., Wöltje, K., Magyar, Z., Pettkó-Szandtner, A., Pélissier, T., De Jaeger, G., Hoth, S., et al. (2020) PP7L is essential for MAIL1-mediated transposable element silencing and primary root growth. Plant J., 102, 703– 717.

