## Supplemental figures for "RTEL1 is required for silencing and epigenome stability"

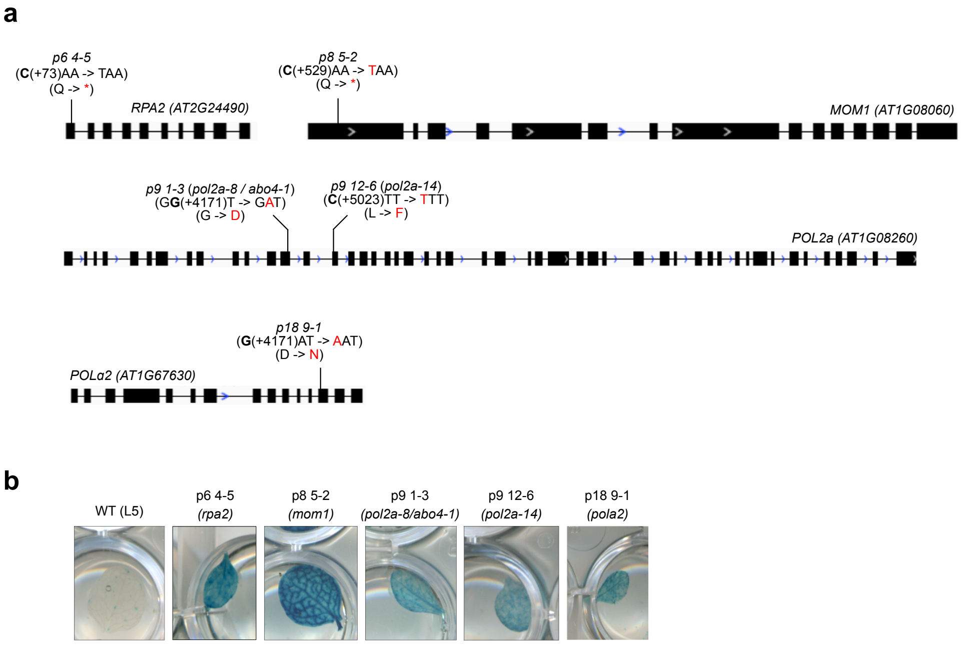


**Supplementary Figure S1: Mutant alleles of known silencing factors isolated in this screen. (a)** Representation of the genes with the mutation and the equivalent amino acid change in red. The numbers in brackets correspond to the position of the mutated nucleotide relative to the transcription start site. **(b)** Photos of the histochemical staining for GUS activity in leaves of the isolated mutants and WT plant (L5) as control.

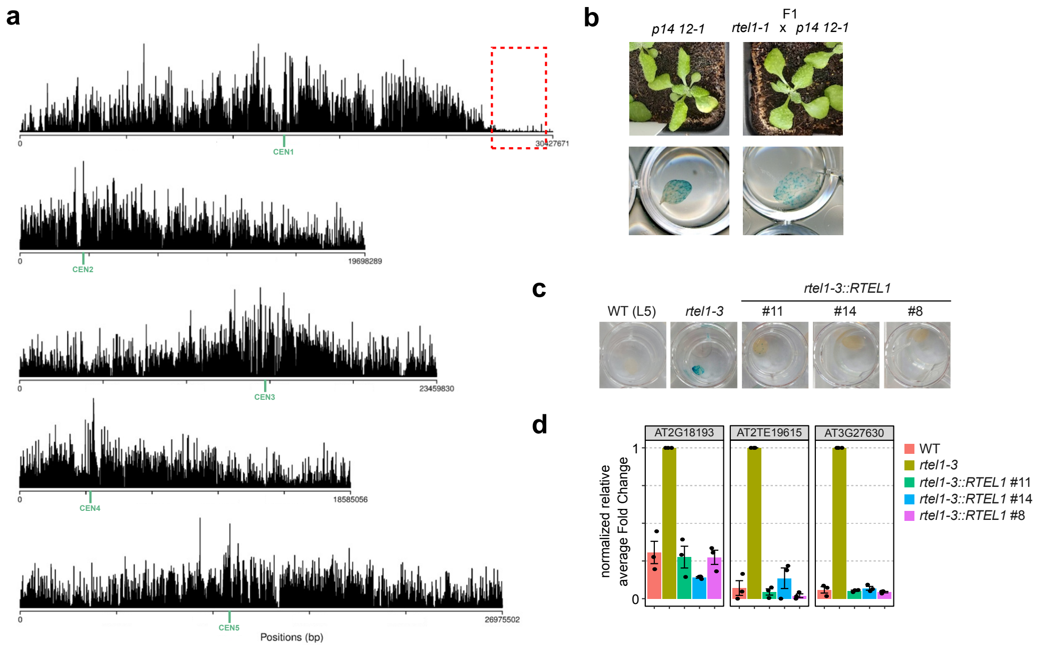


**Supplementary Figure S2: A mutation in *RTEL1* is responsible for silencing defects in *p14 12-1*.** **(a)** Mapping of *p14 12-1* mutation by mapping-by-sequencing Distribution of *Landsberg* (Ler) Single Nucleotide Polymorphism (SNP) frequency along the five Arabidopsis chromosomes in the F2 Ler x *p14 12-1* progeny. The dashed red box marks the region depleted in Ler SNP frequency that carries the causative mutation. **(b)** Photos of 3-week-old plants (top) and histochemical staining for GUS activity (bottom) of the *p14 12-1* mutant and complementation test between *rtel1-1* and *p14 12-1*. **(c)** Representative images of histochemical staining for GUS activity in leaves from plants of the indicated genotypes. Three independent T1 lines were selected for *rtel1-3::RTEL1* complemented genotype. **(d)** RT-qPCR analysis of *rtel1-3* upregulated transcripts from inflorescences of indicated genotyped. Transcript levels are normalized to *ACT2* and further normalized to *rtel1-3* mutant. Sample means are shown with error bars representing standard error of the mean across three technical replicates.

a

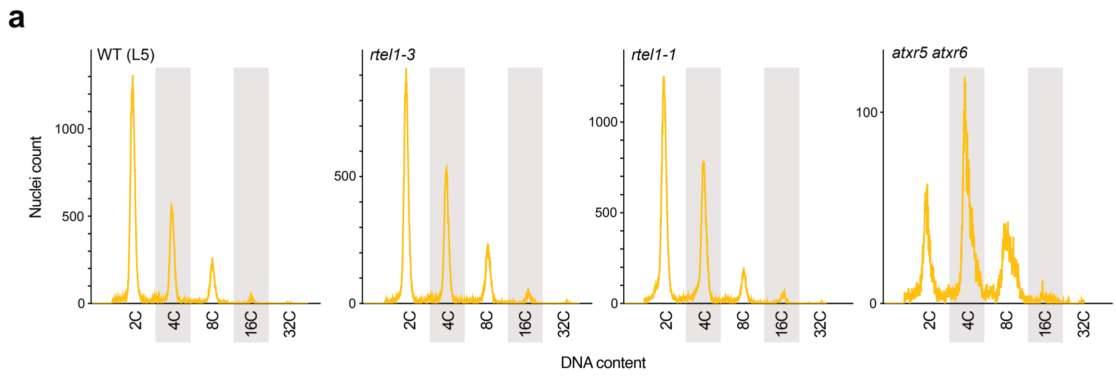


b

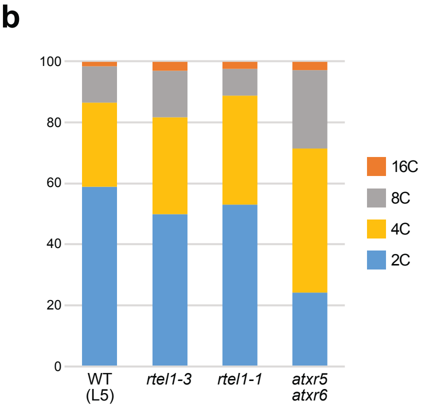


c

| | | L5 | *rtel1-3* | *rtel1-1* | *atxr5 atxr6* |
|---|---|---|---|---|---|
| **2C** | % | **58.9** | **49.8** | **53.0** | **24.2** |
| | Count | 33619 | 22280 | 36834 | 2366 |
| **4C** | % | **27.5** | **31.8** | **35.8** | **47.2** |
| | Count | 15670 | 14228 | 24917 | 4620 |
| **8C** | % | **12.0** | **15.3** | **8.7** | **25.7** |
| | Count | 6873 | 6851 | 6022 | 2517 |
| **16C** | % | **1.4** | **2.8** | **2.3** | **2.7** |
| | Count | 800 | 1259 | 1616 | 267 |
| **32C** | % | **0.2** | **0.2** | **0.2** | **0.2** |
| | Count | 100 | 99 | 115 | 20 |

**Supplementary Figure S3:** DNA content in *rtel1* mutants. **(a)** Flow cytometry profiles of the DNA content of nuclei from the indicated genotypes. **(b)** Comparison of the DNA contents in nuclei of the indicated genotypes. **(c)** Proportions and total nuclei counts with each DNA content for the indicated genotypes.

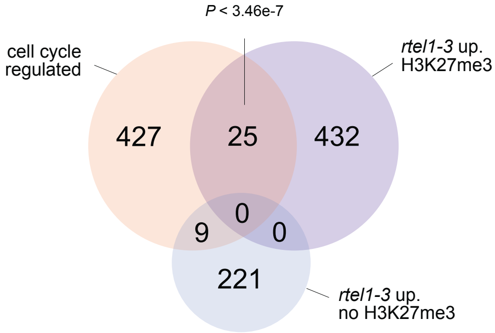


**Supplementary Figure S4:** Venn diagram showing the overlap between *rtel1-3* upregulated PCGs associated on not with H3K27me3 in the WT, and cell cycle-regulated genes (from (1)).

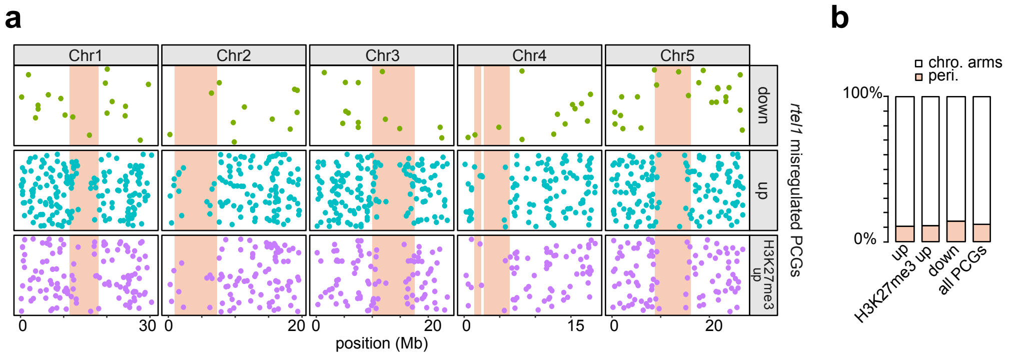


**Supplementary Figure S5: (a)** Genomic localization of *rtel1-3* downregulated PCGs (down), *rtel1-3* upregulated PCGs (up), and *rtel1-3* upregulated PCGs marked by H3K27me3 (H3K27me3 up). **(b)** Proportion of these PCGs located in pericentromeric heterochromatin (peri.) and along chromosome arms (chro. arms).

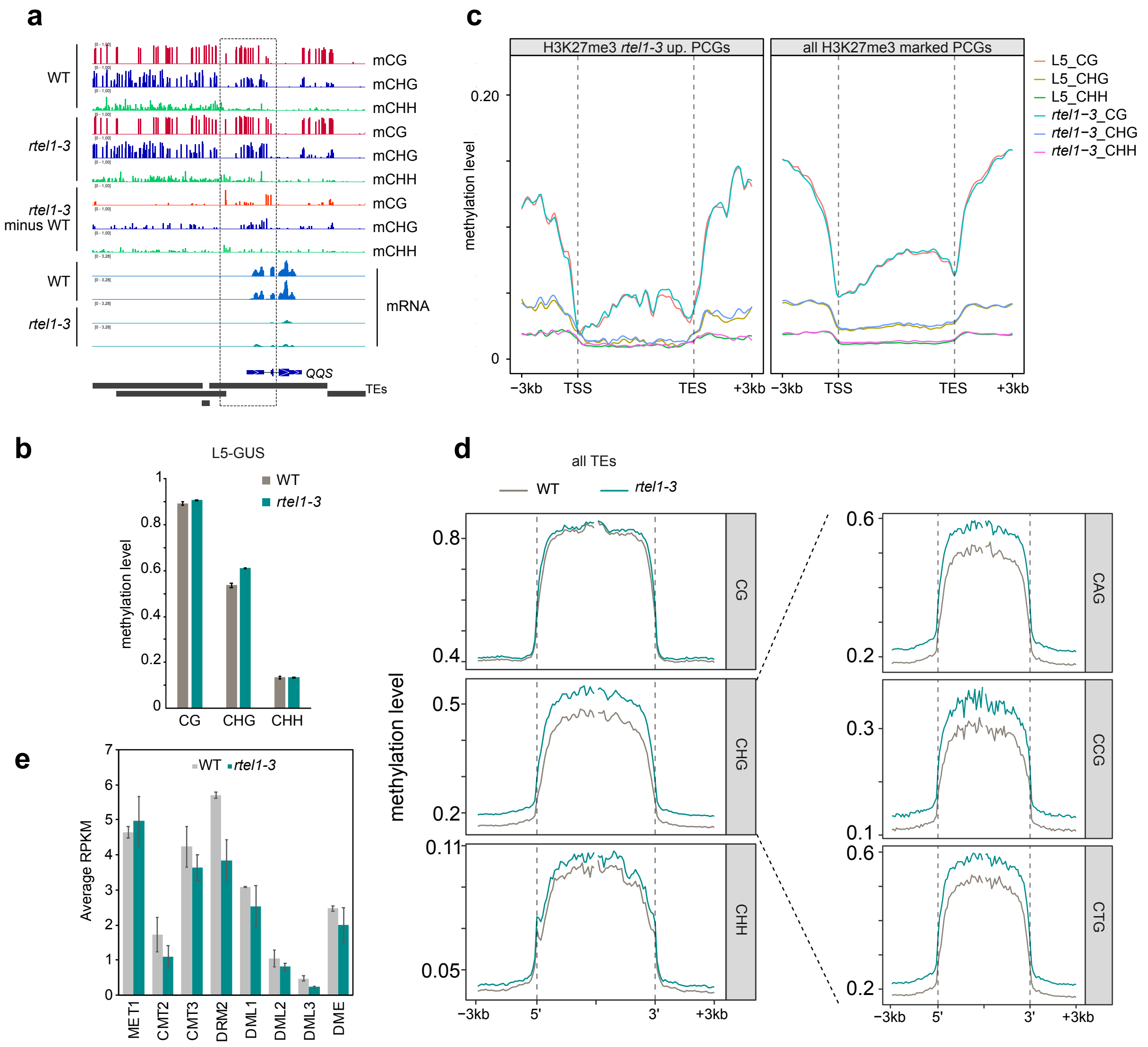


**Supplementary Figure S6: DNA methylation profiles in *rtel1-3* mutant. (a)** Genome browser tracks showing DNA methylation enrichment (in all three cytosine contexts) and mRNA profiles at QQS locus in WT and *rtel1-3*. Local hypermethylation is represented in subtracting WT control WGBS-seq to *rtel1-3*. Two replicates are shown for mRNA-seq data. **(b)** CG, CHG and CHH DNA methylation levels over L5-GUS transgene measured by WGBS-seq in WT and *rtel1-3* plants. Error bars represent standard error of the mean across two biological replicates. **(c)** Metaplots showing average methylation rates in CG, CHG and CHH contexts found in H3K27me3-marked *rtel1-3* upregulated PCGs (left) or in all H3K27me3-marked PCGs of the Arabidopsis genome (right). PCGs were scaled to 6 kb and sequences 3 kb upstream their transcription start site (TSS) or downstream their transcription end site (TES) were included. Average methylation levels per 100 bp windows were plotted. **(d)** CG, CHG, CHH (left) and CHG subcontexts (CAG, CCG and CTG, right) DNA methylation levels over all TEs in WT and *rtel1-3*. TEs were aligned as in Figure 3b. **(e)** Transcript accumulation in reads per kilobase per million mapped reads (RPKM) at the indicated genes for WT and *rtel1-3*. Error bars represent standard error of the mean across two biological replicates.

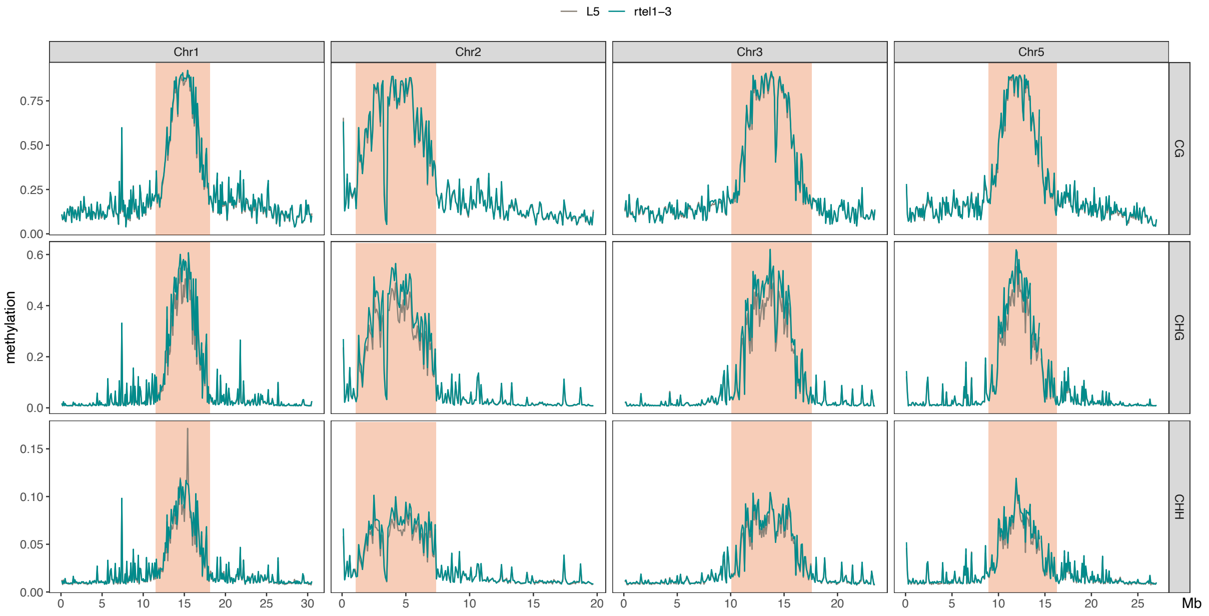


**Supplementary Figure S7:** Average DNA methylation level per 100kb windows on chromosome 1,2,3,5 in CG, CHG and CHH context. Pericentromeric heterochromatin regions are highlighted (coral color). Average DNA methylation levels along chromosome 4 are shown in Figure 3a.

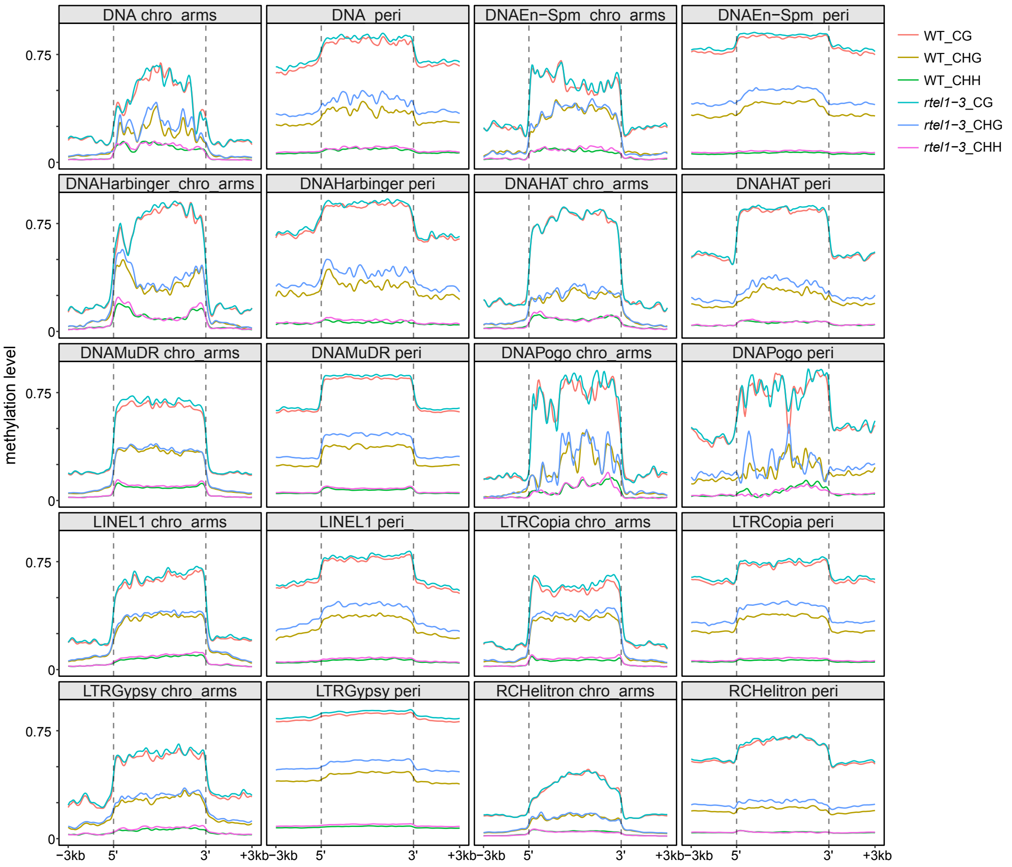


**Supplementary Figure S8:** Metaplots showing average DNA methylation levels in all three cytosine contexts at TE copies from the most prevalent TE superfamilies of the Arabidopsis genome (each with at least 50 copies in pericentromeres and chromosome arms). DNA methylation profiles are shown separately for TE copies located along chromosome arms (chro. arms) and TE copies located in pericentromeric heterochromatin (peri.).

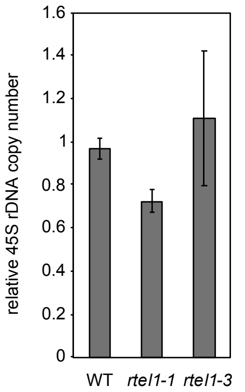


**Supplementary Figure S9: Analysis of rDNA copy number in *rtel1* mutants.** The 45S rDNA copy number was determined by quantitative PCR in *rtel1-1* and *rtel1-3* mutants, in relation to WT (L5 background) plants (n=3). Sample means are shown with error bars representing standard error of the mean across two biological replicates.

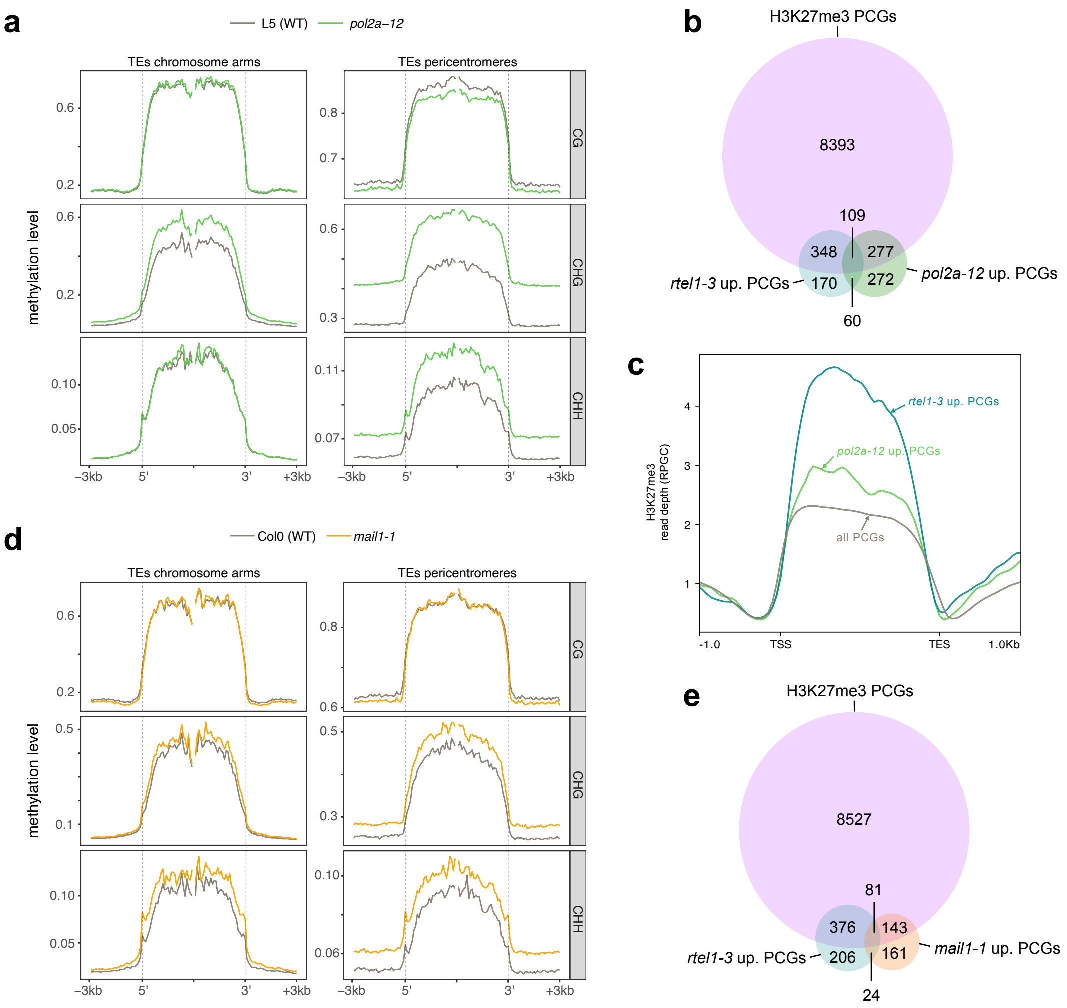


**Supplementary Figure S10: RTEL1 function is mostly independent of POL2A and MAIL1. (a)** Metaplots showing average DNA methylation levels in all three cytosine contexts at TEs located on chromosome arms (left) or in pericentromeric heterochromatin (right) in *pol2a-12* mutant. TEs were aligned as in Figure 3b. **(b)** Venn diagram showing the overlap between *rtel1-3*, *pol2a-12* upregulated PCGs and all H3K27me3-marked genes. **(c)** H3K27me3 ChIP signal over *rtel1-3*, *pol2a-12* upregulated PCGs and all PCGs of the Arabidopsis genome. PCGs were scaled and plotted as in Figure 2d. **(d)** Metaplots showing TE methylation in *mail1-1* mutant represented as in a. **(e)** Venn diagram showing the overlap between *rtel1-3*, *mail1-1* upregulated PCGs and all H3K27me3-marked genes. Methylome and transcriptome of *pol2a-12* and *mail1-1* mutants were re-analyzed from published data (2, 3).

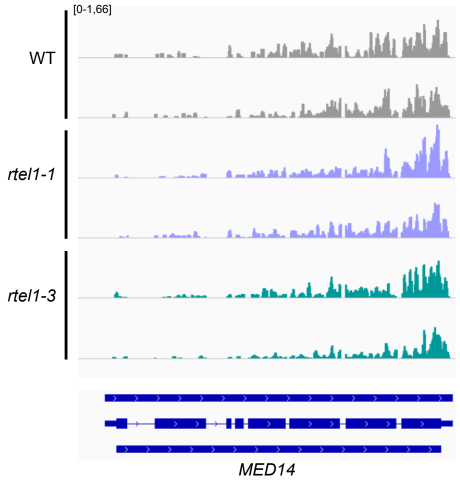


**Supplementary Figure S11: MED14 transcript levels in *rtel1* mutants.** Genome browser tracks showing mRNA profiles at MED14 (AT3G04740) locus in WT, *rtel1-1* and *rtel1-3*. Two replicates are shown.

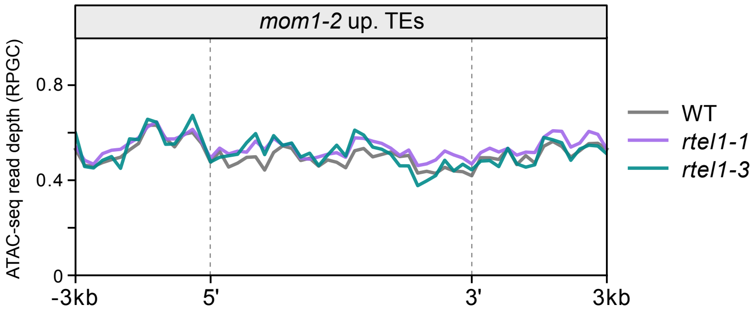


**Supplementary Figure S12: Chromatin accessibility at *mom1-2* upregulated TEs.** ATAC-seq depth over *mom1-2* up-regulated TEs in WT, *rtel1-1* and *rtel1-3* single mutants. TEs were scaled to 6 kb and sequences 3 kb upstream or downstream their 5' and 3' ends respectively were included. Average RPGC signal over 200 bp bins is plotted.

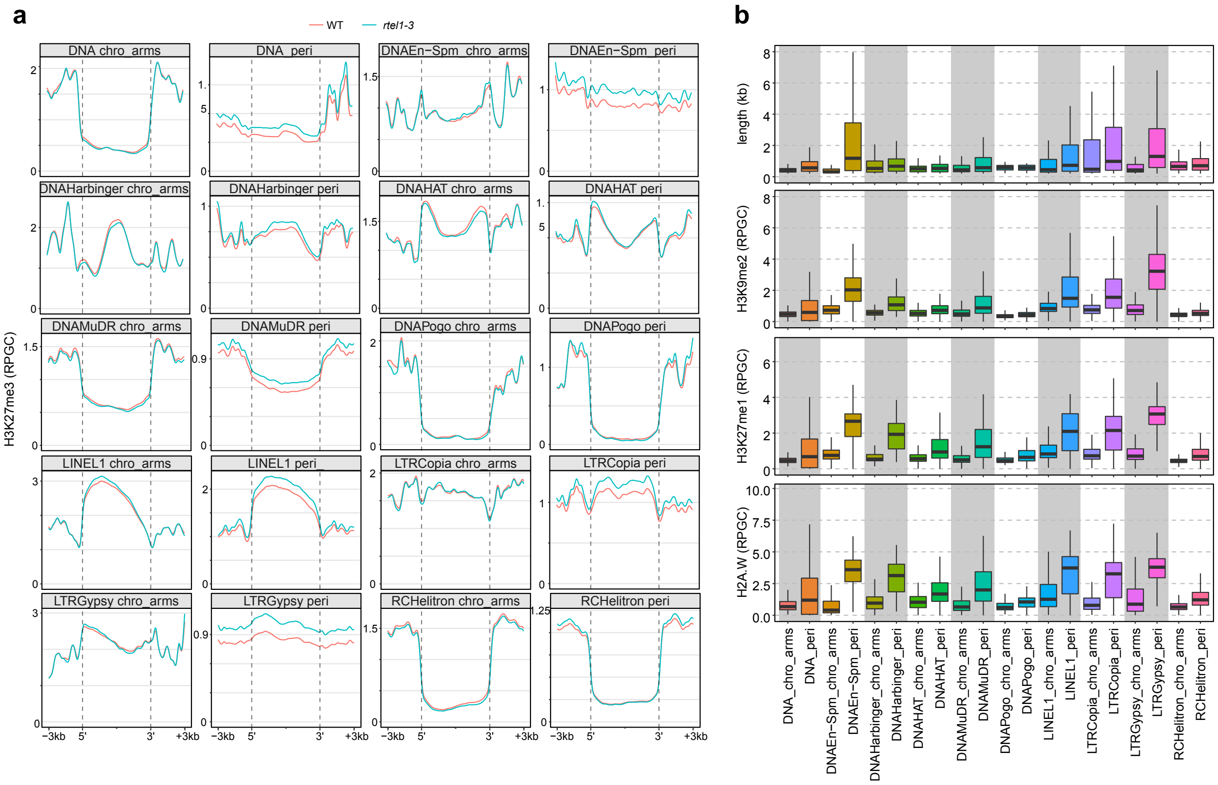


**Supplementary Figure S12: (a)** Metaplots showing average H3K27me3 level at TE copies from the most prevalent TE superfamilies of the Arabidopsis genome (each with at least 50 copies in pericentromeres and chromosome arms). H3K27me3 profiles are shown separately for TE copies located along chromosome arms (chro. arms) and TE copies located in pericentromeric heterochromatin (peri.). **(b)** Boxplot showing length, and ChIP signal for H3K9me2, H3K27me1 and H2A.W at the same TE copies shown in (a).

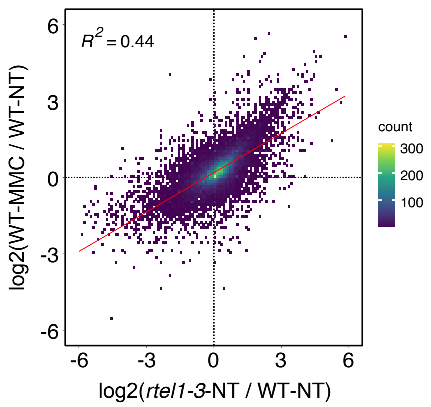


**Supplementary Figure S13: Gene expression changes in MMC-treated WT plants and untreated *rtel1-3* plants.** Scatter plot showing correlation of transcriptional changes of all PCGs in *rtel-3* [log2(*rtel1-3*-NT/L5-NT)] and in MMC-treated WT plants [log2(WT-MMC/WT-NT)]. The colored bin represents the count of PCGs from low (purple) to high (yellow). Linear regression line (red) and $R^2$ ($p < 2.2x10^{-16}$) are indicated.

a

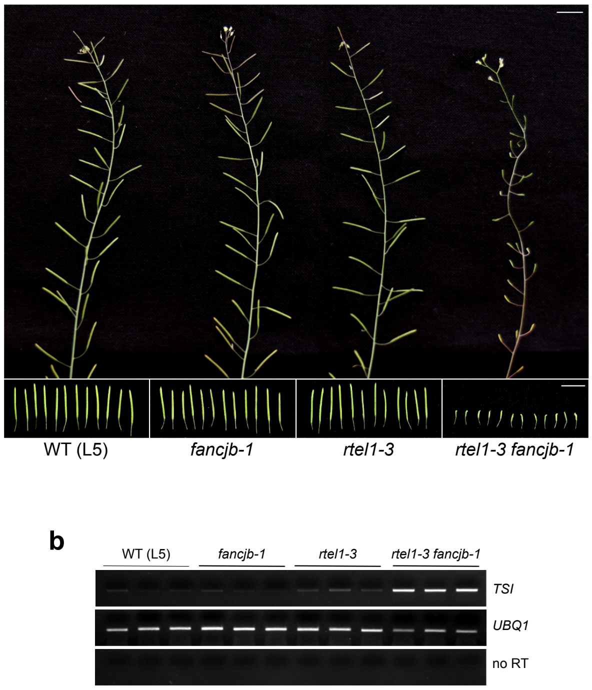


**Supplementary Figure S14: Sterility and silencing release observed in *rtel1-3 fancjb-1* double mutants. (a)** Inflorescence (top) and individual silique (bottom) pictures of the indicated genotype plants. WT (L5 background) plants and single mutants show normal siliques while transformed flowers in *rtel1-3 fancjb-1* double mutants did not self-fertilize and silique elongation did not occur indicating no developing seeds. **(b)** RT-PCR analysis of transcripts from endogenous repeats, TSI (Transcriptionally Silent Information) and 106B. Each line represents biological replicates. Amplification of *UBQ1* (*UBIQUITIN EXTENSION PROTEIN 1*) was used as a loading control. For each target, PCR in the absence of reverse transcription (no RT) was performed to ensure any DNA contamination.

**References:**


1. Menges,M., Hennig,L., Gruissem,W. and Murray,J.A.H. (2002) Cell Cycle-regulated Gene Expression inArabidopsis *. *J. Biol. Chem.*, **277**, 41987–42002.

2. Bourguet,P., López-González,L., Gómez-Zambrano,Á., Pélissier,T., Hesketh,A., Potok,M.E., Pouch-Pélissier,M.-N., Perez,M., Da Ines,O., Latrasse,D., *et al.* (2020) DNA polymerase epsilon is required for heterochromatin maintenance in Arabidopsis. *Genome Biol.*, **21**, 283.

3. Ikeda,Y., Pélissier,T., Bourguet,P., Becker,C., Pouch-Pélissier,M.-N., Pogorelcnik,R., Weingartner,M., Weigel,D., Deragon,J.-M. and Mathieu,O. (2017) Arabidopsis proteins with a transposon-related domain act in gene silencing. *Nat. Commun.*, **8**, 15122.
